# Bioorthogonal photocatalytic quinone methide decaging for cell-cell interaction labeling

**DOI:** 10.1101/2023.04.08.536099

**Authors:** Yan Zhang, Shibo Liu, Fuhu Guo, Shan Qin, Nan Zhou, Xinyuan Fan, Peng R. Chen

## Abstract

Cell-cell interactions (CCIs) play crucial roles in directing diverse biological processes in multicellular organisms, making the high-sensitivity and selectivity characterization of the diverse CCIs in high demand yet still challenging. We herein introduced a bioorthogonal photocatalytic quinone methide decaging-enabled cell-cell interaction labeling strategy (CAT-Cell) for sensitive and spatiotemporally resolved profiling of multitype CCIs. By adapting an optimized quinone methide probe for interacting cell labeling, we demonstrated the excellent capacity of CAT-Cell for capturing CCIs directed by various receptor-ligand pairs (e.g., CD40-CD40L, TCR-pMHC) and further showed its compatibility with tumor-specific targeting systems. Finally, we used CAT-Cell to detect cytotoxic cells (e.g., antigenspecific T cells, Natural Killer cells) in mouse models containing splenocyte mixtures and tumor samples. By leveraging the bioorthogonal photocatalytic decaging chemistry, CAT-Cell offers as a nongenetic, non-invasive and universal toolbox for profiling diverse CCIs under physiological-relevant settings.

## Introduction

Direct cell-cell interactions are critical for diverse cellular activities, orchestrating essential biological processes such as embryonic development and homeostasis^1,2^, highlighting the importance of decoding their interaction specificity under physiological as well as pathological conditions. The development of tumor is closely related to ubiquitous and complicated cellular interactions between cancer cells and tumor infiltrated immune cells co-existed in the tumor microenvironment (TME). Thus, it is crucial to investigate the heterogenous tumor infiltrated immune cell-tumor cell interactions in order to characterize immune responses and tumor progression, and to further guide immunotherapeutic applications.^3-6^ Despite their importance, the highly dynamic nature, spatially restricted distance, as well as complex intercellular environment all made it highly challenging to capture the interacting cells. Furthermore, these interactions in TME can be complicated in terms of compositions and cell types, involving multiple ligand-receptor recognitions with numerous interaction strengths and even the average interfacial gap distances^7^, making systematic characterization of direct interactions of tumor cells more problematic.

Compared to imaging-based methods, proximity labeling approaches have been developed to investigate intercellular interactions by *in situ* tagging within the cellular contact environment, integrating advantages of detection, isolation, and downstream analysis.^8^ For example, enzyme-based strategies have been successfully applied to capture specific interacting cells, but are limited by genetic manipulation or the complicated synthesis of enzyme conjugates.^9-11^ Recently photocatalyst-based approaches have emerged by employing small molecule photocatalysts to generate reactive intermediates, including carbene, singlet oxygen or radical species to drive the proximity labeling process.^12-14^ However, due to the inherent chemical activation mechanisms, chemical tuning of these reactive intermediates is usually difficult, hindering the labeling efficiency or adaptation in complex biological environment. Additionally, these methods are hampered by either the generation of damaging chemical species, increasing the risk of altering cell surfaces and fragile biological environments^15,16^, or the narrow scope of the labeling substrates, which impeded the unbiased detection of multiple ligand-receptor-based cellular interactions. Therefore, a sensitive, non-invasive, and general platform for interacting cell labeling under physiological relevant conditions is still highly desirable but challenging.

Quinone methides (QMs) are a class of unique efficient Michael acceptors that can be targeted to several widely available nucleophilic residues. It can be facially chemical modified for tuning the reactivity, and have been widely used in chemical synthesis and chemical biology.^17-21^ For the inimitable reactivity and controllable rescue property by protection of its phenol oxygen, QMs have become attractive in combination with bioorthogonal decaging chemistry, which has been developed for on-demand activation of bioactive molecules, allowing gain-of-function manipulations via *in situ* rescue of adaptable caging moieties.^22^ In particular, recent bioorthogonal photocatalytic decaging reactions with precise spatiotemporal resolution have shown great potential in widespread applications^23,24^, including proximity labeling of intracellular proteins.^25^ With the flexibility of the quinone methide reactivity, as well as the bioorthogonality and spatiotemporal controllability of the decaging chemistry, we envisaged that photocatalytic decaging of a suitable quinone methide probe could provide a distinct, universal and efficient chemical approach for diverse cell-cell interaction investigations without damaging the fragile biological components.

Herein, we report the development of a photocatalytic quinone methide decaging-enabled cell-cell interaction labeling strategy, termed CAT-Cell, for profiling the highly diverse intercellular interactions (**Fig. 1a**). The iridium photocatalyst was non-genetically equipped on cells to generate reactive QM intermediates upon visible light irradiation and subsequent label targeted diverse protein residues on interacting cells in close proximity. By taking advantage of the chemical tunability of QM probes, our method was sensitive to profile multitype cell-cell interactions within various co-culture systems. Finally, CAT-Cell allowed cytotoxic cells (including antigen-specific T cells and NK cells) identification in primary samples, emphasizing the critical demand of straightforward, non-disturbing and nongenetic operations under living conditions. Together, CAT-Cell provides a valuable chemical approach for sensitive and universal profiling of multiple cell-cell interactions even without prior knowledge of molecule pairs in diverse living systems.

**Figure 1.**
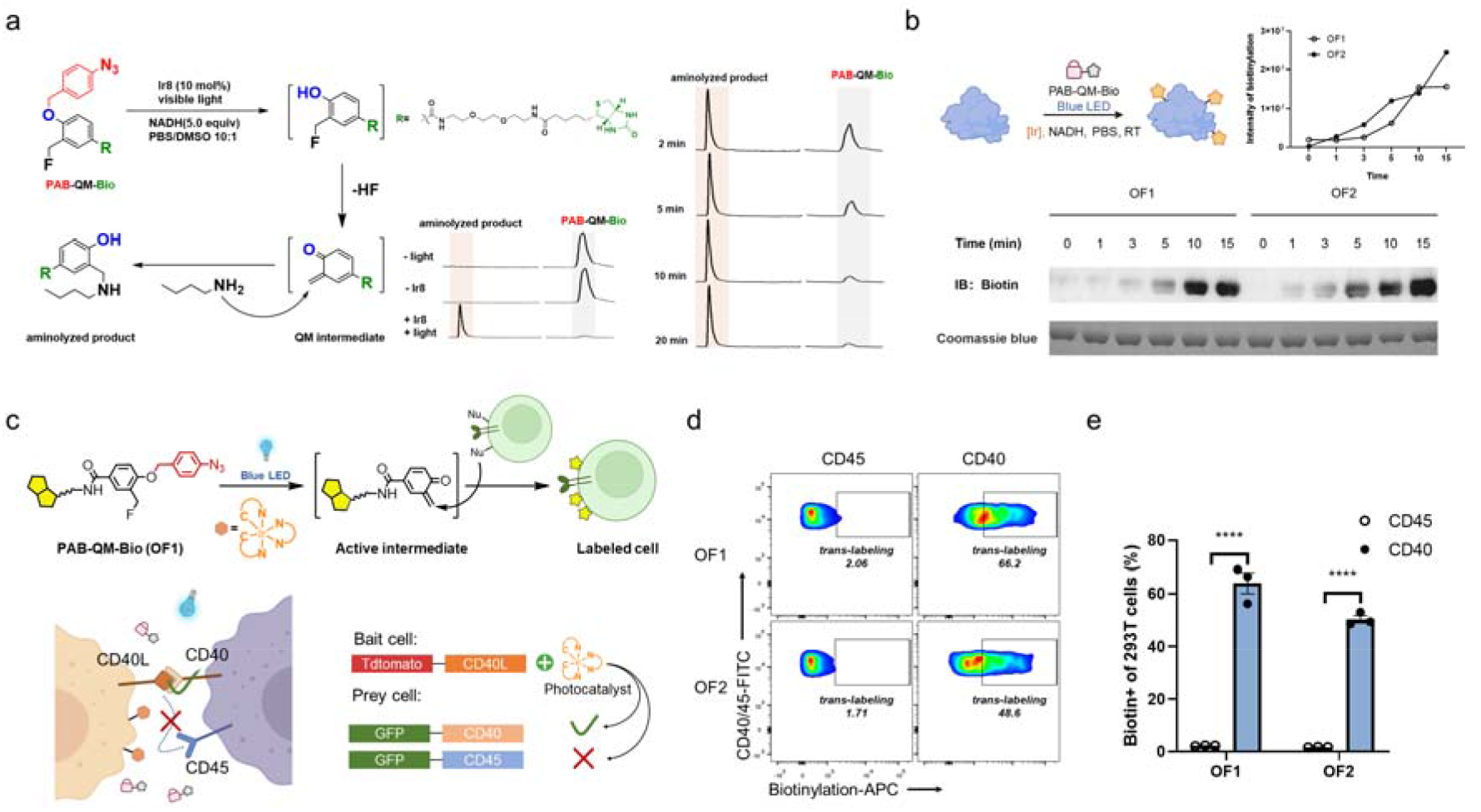
Design and development of bioorthogonal photocatalytic quinone methide decaging-enabled cell-cell interaction labeling (CAT-Cell). (a) Design of quinone methide-based labeling probe and HPLC traces of photocatalytic decaging of PAB-QM-Bio (OF1) in aqueous media. HPLC traces of the photocatalytic decaging of PAB-QM-Bio (OF1) probe under blue LED at given time points. Conditions: 0.1 mM PAB-QM-Bio (OF1), 100 mM butylamine, 0.5 mM NADH, 10 mol % Ir8, in PBS/DMSO (10/1) buffer, irradiated by blue LED (4 mW/cm^2^) for 20 min. (b) *In vitro* evaluation of different QM probes by CAT-Cell enabled BSA labeling at given time points. Biotinylated BSA and total BSA were analyzed by immunoblotting and Coomassie blue staining, respectively. (c) Experimental setup to analyze intercellular labeling in transfected HEK293T cells by CAT-Cell. Photocatalytic iridium compound was conjugated on the cell surface via N-hydroxysuccinimide ester– based lysine coupling reaction, allowing bioorthogonal decaging reaction-enabled biotinylation by well-designed caged PAB-QM-Bio (OF1) probe for cells in close proximity under mild blue light irradiation. The CD45-positive HEK293T cells were set to be the control group compared to CD40positive cells. (d, e) Flow cytometric analysis (f) and bar graphs (g) showing different biotinylation efficiencies of QM probes. Data are presented as mean ± SEM (n=3); ns, p > 0.05; *p < 0.05; **p < 0.01; ***p < 0.001; ****p < 0.0001; two-way ANOVA followed by Sidak’s multiple comparisons test. Artworks of (b) and (c) were created with BioRender.com.

## Results and discussion

### Development and validation of photocatalytic QM-decaging for CCI labeling

We started by developing an intercellular labeling system based on quinone methide chemistry via photocatalytic decaging. In our previous intracellular protein labeling system, high reactive OF2 quinone methide was used for subcellular proteomics profiling.^25,26^ However, given the spatially restricted distance and complex intercellular environment, larger labeling radius of quinone methide ensuring sufficient labeling efficiency was needed, which in turn required quinone methide with moderate reactivity resulting from slower quenching rate in aqueous environment. Inspired by the feasible chemical tunability of the quinone methide chemistry, we designed the OF1 probe. Due to the loss of the electron-withdrawing fluorine group comparing with the OF2 counterpart, the quinone methide intermediate of OF1 is expected to be quenched slower because of its less electrophilic reactivity to the nucleophiles including water in the aqueous environment. The photocatalytic decaging of the PAB-protected OF1 probes in aqueous solution were monitored using HPLC first (**Fig. 1a**). Slowly conversion of OF1 into the hydrolyzed product was detected in the presence of both Ir8 (10 mol %) and blue LED irradiation. Nucleophilic addition product was obtained when n-butylamine was added as nucleophile (**Fig. 1a**). Given that labeling reactions on cells usually occur on surface proteins, we next evaluated the protein labeling ability taking BSA as a model protein and the OF2 probe as the reference. As expected, slower labeling rate for OF1 was detected due to its less electrophilic reactivity (**Fig. 1b**).

Next, we aimed to evaluate the capability of intercellular labeling by quinone methide probes, and cell-cell interactions via the CD40-CD40L pair, an important ligand-receptor pair influencing immune activation or suppression^27^, was designed and constructed as a model assay (**Fig. 1c**). Two groups of HEK293T cells were transfected with CD40 and CD40L, respectively. Then the CD40L+ cells were equipped with iridium photocatalyst to form the bait cells by treating with NHS-[Ir] (**Fig. S1b**). Flow cytometry analysis showed significant biotin signal on the cell surface. We further confirmed membrane protein labeling via western blot analysis, and chose the 100 µM as optimal concentration of NHS-[Ir] for labeling (**Fig. S1c, d**). The CD40+ cells were then co-cultured as prey cells and treated with PAB-QM-Bio probes (1 µM), NADH (500 µM), and 5 minutes of blue LED irradiation. After optimization of irradiation, we were delighted to observe significant biotin signals on the CD40+ prey cells via flow cytometry analysis. In contrast, only trace labeling was observed when using CD45+ cells as a non-interacting control (**Fig. 1d, e**), confirming the labeling specificity of which simple colocalization between non-interacting cells and bait cells within the same microenvironment was not adequate to drive labeling process. Additionally, due to the more suitable labeling radius of the less reactive OF1 intermediate, it demonstrated higher labeling efficiency than the OF2 probe (63.9% versus 50.3% labeling ratio) in the trans-cell labeling, as expected. These results confirmed the capability of photocatalytic decaging chemistry for specific capturing of the cell-cell interactions, as well as the tunability and flexibility of the labeling probes by decaging strategy.

Besides, as a visible light-triggered reaction, the precise photo switch through easily tunable parameters such as light intensity or duration of CAT-Cell offered a convenient way to achieve spatial specificity when capturing interacting cells in precise regions. As a proof of concept, we cocultured HEK293T cells transfected with CD40 and CD40L separately and CD40L+ cells were pretreated with [Ir]. Then we chose a small region of interest (ROI) for photo irradiation, resulting in CAT-Cell-introduced biotinylation (**Fig. S2a**). The labeling efficiency of each well was represented as a “pixel” in the heatmap, in which biotinylation was successfully induced when spatial selectivity to the ROI and cell-cell interaction specificity to the CD40+ cells were simultaneously achieved (**Fig. S2b, c**). The spotless background in the dark region minimized interference from other incoherent zones in complex samples, facilitating the analysis of specific cell-cell interactions.

### CAT-Cell is a sensitive, modular and targetable CCI labeling strategy

With the established CAT-Cell platform in hand, we sought to explore the versatility of this method for various situations especially for identification of more delicate interaction pairs.

There are multiple ligand-receptor-based cellular interactions in complex systems, and T cell receptor (TCR)-cognate peptide-major histocompatibility complex (pMHC) recognitions are crucial for orchestrating adaptive immunity, regulating immune surveillance and homeostasis with the help of other adhesion molecules and co-stimulatory/checkpoint receptor-ligand pairs.^28-30^ However, it is still challenging to detect specific TCR-pMHC interactions due to the highly diversity and low affinity of TCR repertoires. Thus, we further explored the labeling efficiency and specificity of CAT-Cell in this delicate interaction model. NY-ESO-1 is one of the most immunogenic tumor antigens, widely expressed in various cancers, and is considered an ideal target antigen for tumor immuno-therapy.^31,32^ K562 cells expressing HLA-A2:02*01 could present the NY-ESO-1157-165 antigen peptide, which can be specifically recognized by JC5 T cells expressing 1G4 TCR but no endogenous TCR. NY-ESO-1-3Y (3Y) peptide-primed K562 cells were modified by NHS-[Ir] and co-cultured with 1G4 TCR JC5 cells, followed by CAT-Cell-based biotinylation under blue LED irradiation. Considering the differences in intensity and composition of the CD40-CD40L and TCR-pMHC interactions, we benchmarked the labeling performance of OF1 against OF2 once more. As expected, OF1 showed much more evident labeling efficiency than OF2 (36.3% versus 12.6%), confirming the activity difference between the two probes. Significant biotin signals were detected with the 3Y priming group (36.3%) compared to the no peptide group control (8.9%), indicating the achievement and high specificity of CAT-Cell in a delicate CCI system (**Fig. 2d, e and Fig. S3**).

**Figure 2.**
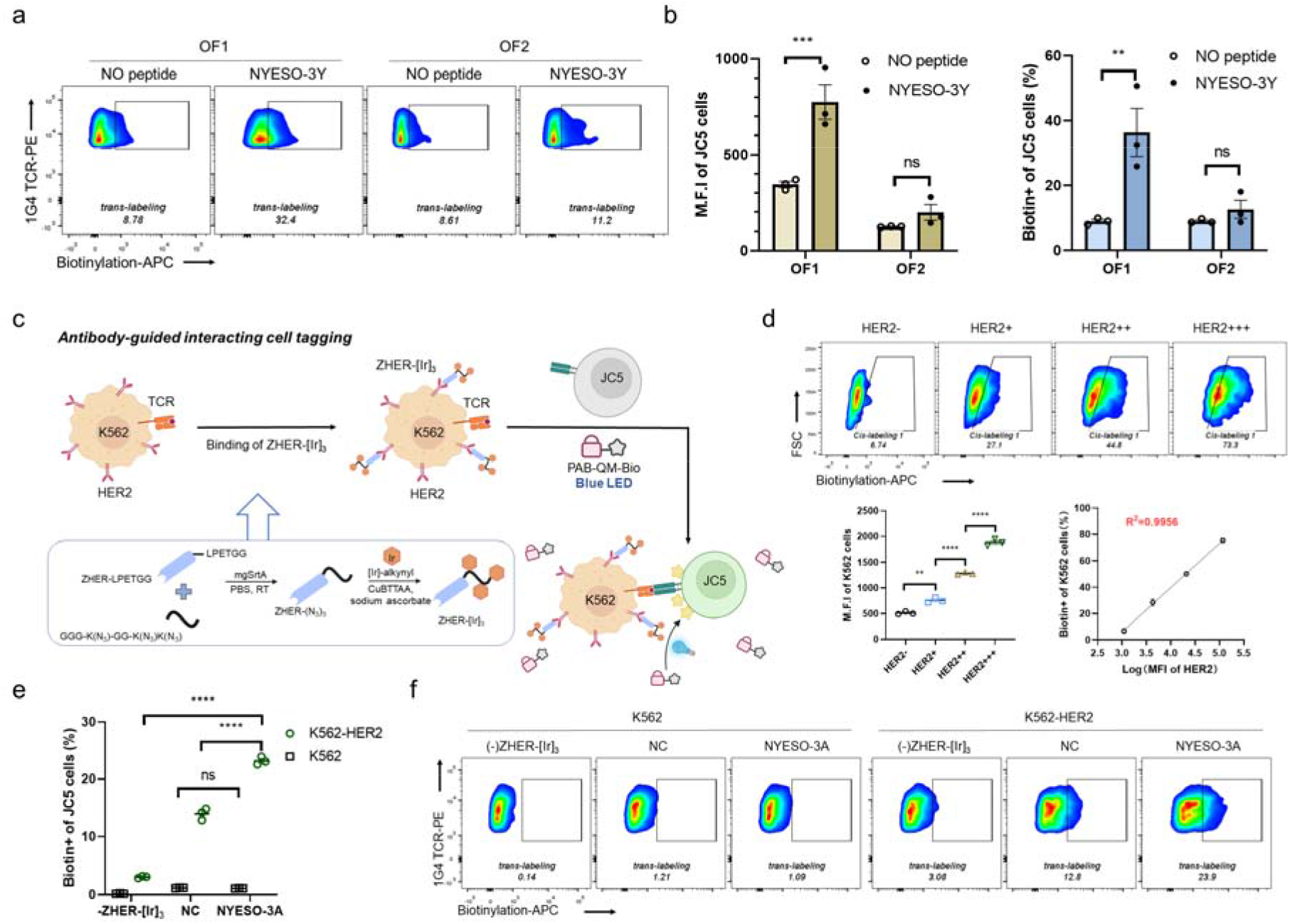
CAT-Cell enables sensitive, modular and targetable labeling of CCIs. (a, b) Flow cytometric analysis of CAT-Cell for detecting TCR-pMHC pair based-CCIs. The results showed significant efficiency difference between QM probes for antigen-specific biotinylation of pMHC-TCR pair induced-CCIs. HLA-A2:02*01 K562 cells not primed with any antigen peptide (“no peptide” group) was set as negative control. (c) Schematic view of intercellular labeling via CAT-Cell in combination with nanobody-photocatalyst conjugate (ZHER-[Ir]_3_). [Ir] was conjugated with ZHER via mgSrtA-mediated ligation and click reaction. Created with BioRender.com. (d) ZHER-[Ir]_3_ binding to target cells introduced significant *cis*-labeling under blue light irradiation, and biotinylation efficiency increased along with HER2 expression. R square=0.9956. (e, f) Flow cytometric analysis of pMHC-TCR interaction-based biotinylation introduced by ZHER-[Ir]_3_ enabled CAT-Cell. The biotin signals in control groups in which K562 cell without HER2 were all at background level.

Next, we aimed to investigate the integration of our CAT-Cell with a targeted system that could direct [Ir] localization to a specific cell population. To achieve this, we constructed a nanobody-photocatalyst conjugate using our previously evolved mono-glycine recognition Sortase A (mgSrtA) and CuAAC click reaction.^9^ This yielded ZHER-[Ir]_3_ conjugates with three site-specific photocatalysts per ZHER for targeted binding to HER2+ cells (**Figure 2c and S4a**). ZHER-[Ir]_3_ was used to bind with K562 cells, followed by employment of CAT-Cell (**Figure 2c**). The HER2+ K562 cells were divided into four groups based on the expression level of HER2, in which the *cis*-labeling ratios were positively correlated with the abundance of HER2 (**Figure 2d**). We also examined *trans*-labeling in NY-ESO-1-3A (3A) peptide-induced CCI system and took the negative control peptide (NC) as a comparison, which binds to HLA-A2:0201 but has no interaction with 1G4 TCR. Significant biotin signals on the HER2+ cells compared to negligible signals on the HER2-cells, as well as the noteworthy biotinylation difference between 3A and NC group, demonstrated the high selectivity and specificity of our targeted strategy in detecting cell type-specific interacting cells (**Figure 2e, f**). Additionally, we observed similar high specificity and efficiency of targeted labeling in the CD40-CD40L system (**Figure S4b-d**).

### *Ex Vivo* profiling of antigen-specific T cells and NK cells

Considering CAT-Cell’s reliability in detecting antigen-specific JC5 T cells with high accuracy, and the unbiased nature of the decaging reaction-based labeling method, we next assessed its feasibility in primary tissue samples for identification interacting cytotoxic immune cells. We chose the model antigen ovalbumin (OVA)-primed MC38 murine colon carcinoma cells to label CD8+ T cells in OT-I transgenic mice splenocytes. We co-cultured splenocytes with MC38 cells treated with NHS-[Ir] and different antigen peptides, followed by irradiation with blue LED light in the presence of the OF1 probe (**Figure 3a**). We detected an increased biotin signal and T cell activation marker CD69 signal with the OVA_257-264_ treated group compared to the GP_33-41_ group or no peptide control, and the two signals correlated with each other (**Figure 3b,c and S5a, b**). In addition to CD8+ T cells, natural killer (NK) cells are also important cytotoxic lymphocytes targeted tumor cells to inhibit its development, and recently, there’re growing enthusiasm in developing NK cell therapies’ potential.^33,34^ Thus, we also evaluated the labeling efficiency of CAT-Cell to capture NK cells. We used the MC38-B2M KO cell line which lacks mature MHC-I structure and MC38-WT cell line as the bait cells and employed CAT-Cell to profile the difference of interactions between NK cells and target cells (**Figure 3a**). We observed obvious biotin signals on NK cells in both groups, confirming labeling capacity of CAT-Cell for NK cells. What’s more, there was increased biotinylation on the NK cells co-cultured with MC38-B2M KO rather than normal MC38 cells, demonstrating enhanced identification and interaction of NK cells and tumor cells due to the lack of MHC-I (**Figure 3d and S5c**).

**Figure 3.**
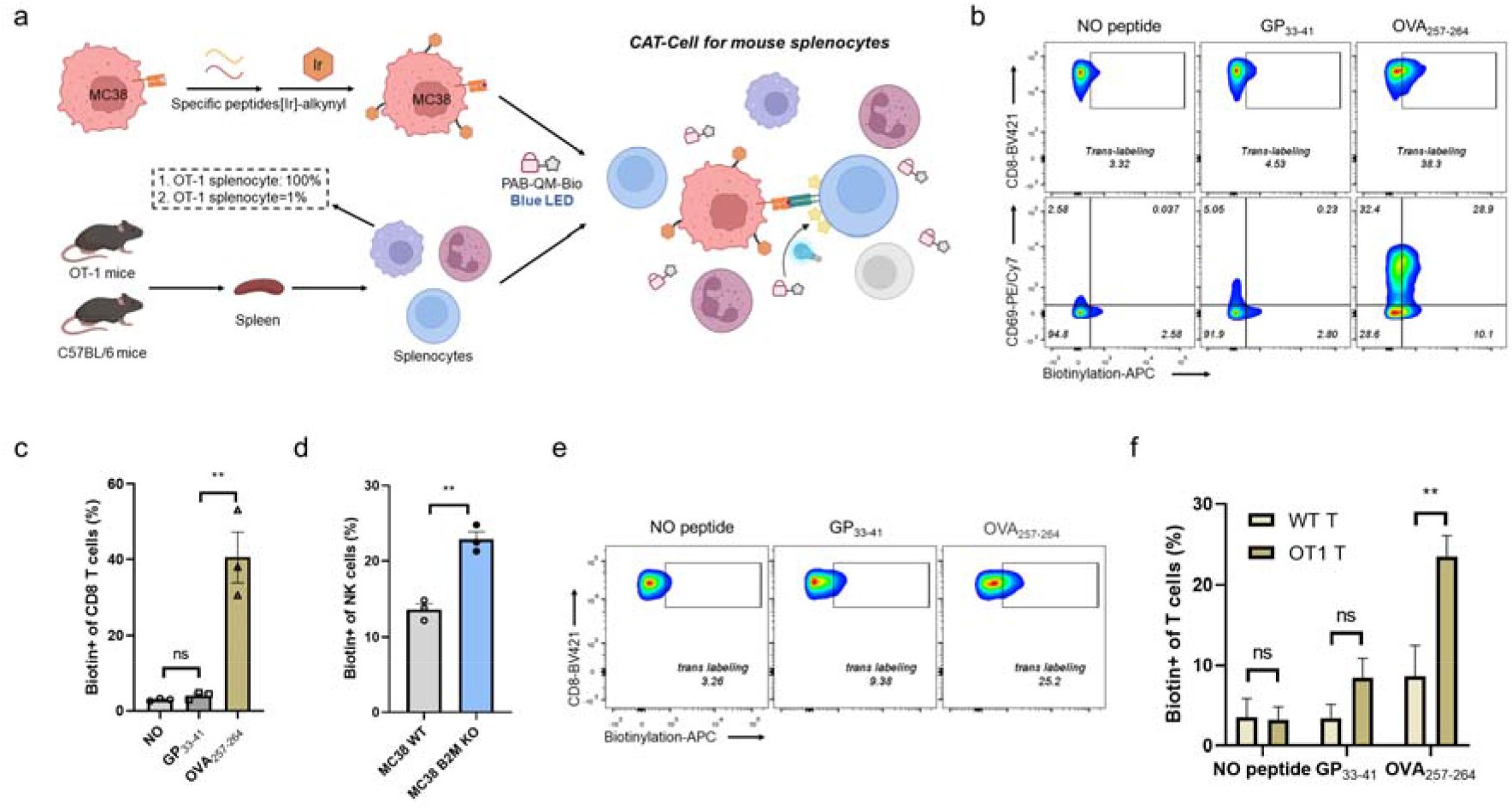
CAT-Cell enabled *ex vivo* identification of antigen-specific T cells and NK cells in mouse model. (a) experimental setup for CAT-Cell enabled *ex vivo* profiling of antigen-specific T cells and NK cells in mouse model. OT-I splenocytes were labeled by antigen-primed [Ir]-MC38 or MC38 B2M KO cells after 2 h incubation. OT-I T cells can recognize and interact with MC38 through the specific interaction between transgenic TCRs and OVA_257−264_ peptide-loaded MHC molecules while T cells from C57BL/6 mouse cannot. And the interaction between MC38 B2M KO cells and NK cells can be strengthened due to the loss of mature MHC complex compared with WT MC38 cells. (b, c) Representative flow cytometric analysis of antigen-specific T cell biotinylation in OT-I splenocytes. GP_33▫41_ (KAVYNFATM) was a nonspecific antigen peptide from the lymphocytic choriomeningitis virus (LCMV). The biotin signals in OVA_257−264_ groups correlated with GFP signals in (b). (d) Flow cytometric analysis of primary NK cells interacting with different MC38 cells. The loss of mature MHC structure made MC38 B2M KO cell easier to active NK cells so that the interaction can be enhanced as well as the biotin signals. (e, f) Characterization of sensitivity and specificity of CAT-Cell in multicellular environment. Splenocytes from OT-I mouse and C57BL/6 mouse were mixed up to mimic a complex multicellular environment, which followed by interacting cell profiling via CAT-Cell. Flow cytometric analysis of antigen-specific T cell biotinylation of splenocytes. Only the OT-I T cells from OT-I mouse can be biotinylated specifically.

To investigate the selectivity and specificity of CAT-Cell, we constructed a mixture system consisting of splenocytes from OT-I mice and C57BL/6 mice and employed CAT-Cell in the mimic system (**Figure 3a**). We detected increased biotin signals only on the OT-I T cells of the OVA_257-264_ group, as well as activation signals (**Figure 3e, f and S5d, e**). In contrast, an insignificant labeling background was observed on WT T cells of the OVA_257-264_ group or OT-I T cells of the GP_33-41_ group, likely due to the “non-antigen-specific” interactions existing beyond pMHC-TCR on the cell surface.

### Identification of antigen-specific T cells in tumor-infiltrating lymphocytes

The success of applying CAT-Cell to capture antigen-specific T cells in pure or mixed splenocytes prompted us to interrogate the interactions of tumor with tumor-infiltrating lymphocytes (TILs) and further investigate the probability of profiling the composition of TILs in TME. We employed MC38-OVA and MC38-B2M KO cell line as models. These two kinds of solid tumors were digested into single-cell suspensions, and MC38 cells with [Ir] pretreatment were respectively loaded with OVA_257-264_ or GP_33-41_ peptide as different bait cells to co-culture with these tumor single-cell suspensions, followed by the interacting TILs labeling via CAT-Cell (**Figure 4a**). To our surprise, only the incubation of OVA_257-264_ primed MC38 cells and MC38-OVA tumor single-cell suspensions led to the significant biotinylation signals of CD8+ T cells, indicating the primary antigen-specific CD8+ T cells could be detected successfully (**Figure 4b,c**). By contrast, there was minimal *trans*-labeling of CD8+ T cells when GP_33-41_ primed MC38 co-cultured with MC38-OVA tumor single-cell suspensions, suggesting the specificity of the labeling method. Moreover, only background-level biotinylations were detected in both groups involving MC38-B2M KO tumor single-cell suspensions, likely due to the absence of mature MHC complexes, which impacts the antigen presentation as well as the antigen-specific T cells infiltrating. Finally, we characterized the molecular phenotypes of Biotin+ and Biotin− CD8+ T cells, and found the Biotin+ CD8+ T cells had higher expressions of PD-1, TIM-3, CD39 and CD137 (**Fig. 4d and S6**), indicating that interacting CD8+ T cells exhibiting an activation and/or dysfunction phenotype.^35-40^ Together, these results indicated that CAT-Cell was competent to identify antigen-specific T cells with high selectivity and susceptivity in natural complex multicellular environment, and showed the potential to profile unique component difference of TME.

**Figure 4.**
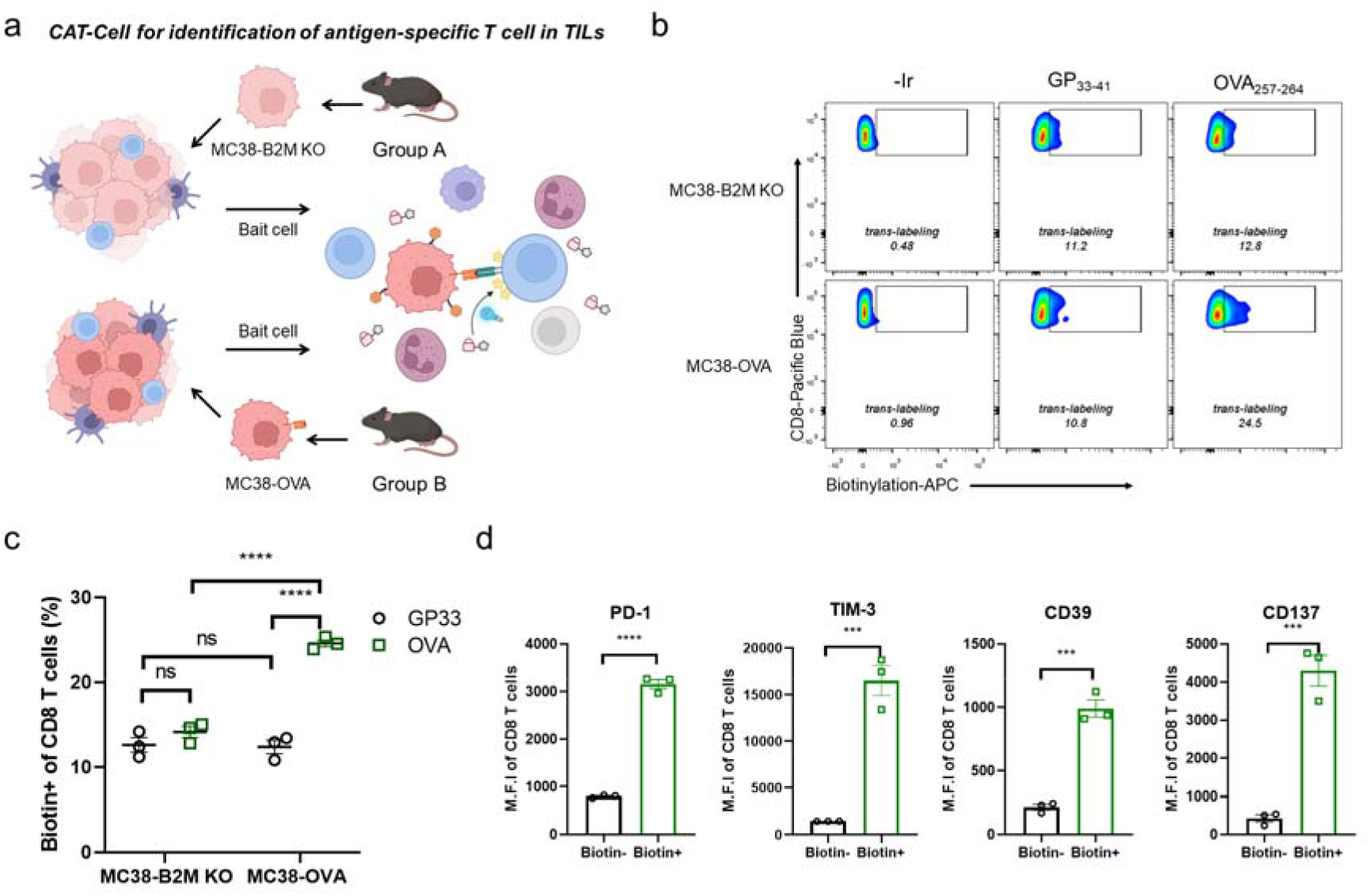
Identification of antigen-specific T cells in tumor-infiltrating lymphocytes by CAT-Cell. (a) Schematic illustration of antigen-specific T cells identification via CAT-Cell in MC38 tumor models. The C57BL/6 mice were planted with MC38 or MC38 B2M KO cells separately and harvested subcutaneous solid tumors were digested into single-cell suspensions. After tumor-T interaction, CAT-Cell-induced labeling was carried out, and the biotin signal was analyzed by FC. Created with BioRender.com. (b, c) Flow cytometric analysis and summary statistics of antigen-specific biotinylation of TILs in MC38 tumor models, n = 3. (d) The expression intensities of potential functional markers on biotin+/− CD8+ T cells according to flow cytometric analysis. n = 3.

## Conclusion

In summary, we have developed a photocatalytic QM-decaging system that enabled intercellular interaction labeling via the bioorthogonal decaging chemistry. Compared to existing photocatalysis-enabled cell capturing methods, CAT-Cell employed distinct QM labeling chemistry, which showed unique advantages. First, the inimitable reactivity of QMs for widespread nucleophilic protein residues cross multiple cell types made its sensitivity sufficient for detecting the weak and highly dynamic intercellular interactions in an unbiased manner regardless of ligand-receptor involved. Secondly, the chemical tunability of QMs offered a facial approach to improve the labeling efficiency ensuring its extensive application in complex biological scenarios. Additionally, the bioorthogonal QM decaging strategy was more biologically benign and non-invasive, avoiding the potent damaging of prey cells by reactive species previously used for proximity cell labeling such as singlet oxygen, radicals or peroxides. Finally, by coupling the QM chemistry with easily prepared iridium photocatalyst, CAT-Cell is straightforward, non-genetic and with high spatiotemporal resolution, which is highly suitable for primary samples under *ex vivo* or even *in vivo* conditions.

In this work, we designed and developed CAT-Cell to capture multiple ligand-receptor-induced cellcell interactions and showed its capacity in both cultured cells and primary samples. The sensitivity and selectivity of the labeling method were demonstrated in the TCR-pMHC system and the broad utility was proven by detection of multiple cell types (including HEK293T, antigen-specific T cells and NK cells). By rational design and establishment of quinone methide probes and platforms, we provided a photocatalytic decaging method differing from those doublet-based methods requiring external computational assistant^41^ or contact-dependent methods which may have concerns due to the artificial cellular interaction interference.^42,43^ It is also complementary to the proximity labeling methods based on specific enzyme or chemical intermediates for intercellular interaction labeling.Considering the chemical tunability of caged QM probes, we expect that there will be more new designs fitting other application scenarios or further improving the labeling efficiency for cell-cell interaction profiling. The design of CAT-Cell is also modular and programmable, permitting the wide adaptability for broad applications. We are currently coupling the CAT-Cell strategy with downstream analyses such as single-cell RNA sequencing, TCR sequencing, and proteomics profiling of tagged cells, which may shed light on the mechanisms of crucial cellular interactions in the immunological process as well as immunotherapy treatment from clinical samples.

**Scheme 1.**
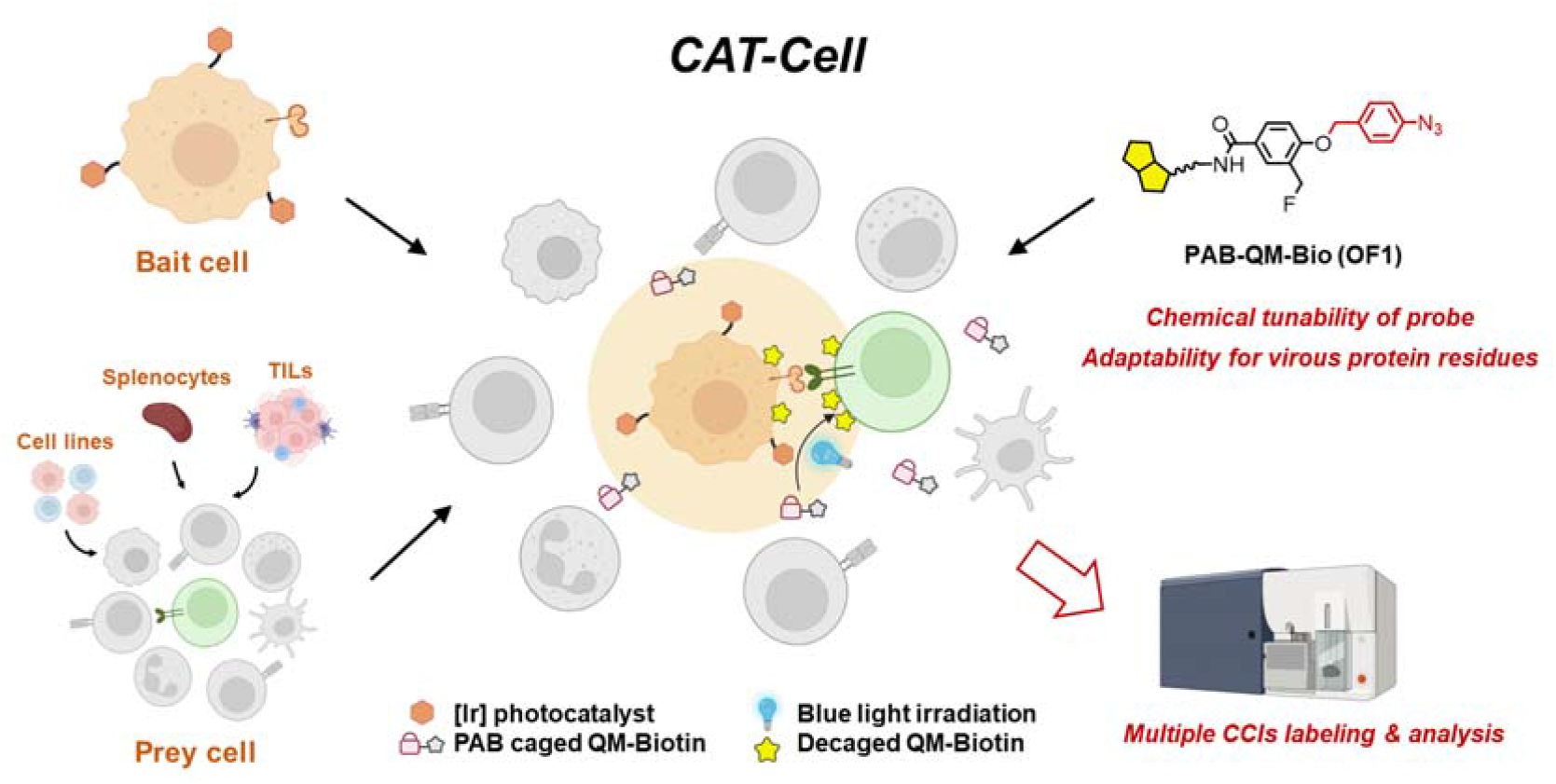
Bioorthogonal photocatalytic quinone methide decaging-enabled multiple cell-cell interaction labeling.

## Associated content

### Supplementary information

Materials, protocols and data characterizations for all biological and chemical experiments are described in detail in the Supplementary information.

### Author information

P.R.C., X.F. and S.L. conceived the study. X.F. and P.R.C. supervised the project. Y.Z., F.G., S.Q. and N.Z. performed the experiments and analyzed the data. S.L. helped with data analysis, project discussion and manuscript preparation. Y.Z., X.F. and P.R.C. wrote the paper with input from all authors.

### Notes

The authors declare no competing interests.

## Acknowledgment

We acknowledge funding from the National Natural Science Foundation of China (22222701, 22077004, 92253301, 21937001, 22137001), the Ministry of Science and Technology (2019YFA0904201, 2022YFA1304700, 2022YFE0114900), Beijing Natural Science Foundation (Z200010) and Li Ge-Zhao Ning Life Science Junior Research Fellowship.

## Notes

### Competing Interest Statement

The authors have declared no competing interest.

### Summary of Updates

Section on TILs identification updated to characterize the phenotype of interacting CD8 T cells; author affiliations updated; Supplemental files updated.

