## Supplementary information for "Bioorthogonal photocatalytic quinone methide decaging for cell-cell interaction labeling"

**Table of Contents**

**Material and Methods**

**1. Chemistry**

General considerations

Synthesis of MNP-caged QM probe

Synthesis of PAB-caged QM probe

Synthesis of photocatalysts (Ir_3_-PEG_4_-NHS)

Synthesis of nanobody-photocatalyst conjugate (ZHER-Ir_3_)

HPLC analysis of photocatalytic decaging of PAB-QM-Bio

Photocatalytic decaging of PAB-QM-Bio for BSA labeling

**2. Biology**

General considerations

Cell line culture and Mice

Primary cells preparation and culture

Photocatalytic proximity labeling of PAB-QM probes in CD40L-CD40 system

Photocatalytic proximity labeling in pMHC-TCR system

CAT-Cell protocol for OT-I antigen-specific T and NK cell capture

CAT-Cell protocol for detection of antigen-specific T cells in tumor samples

**Supplementary Figures**

**References**

**NMR spectra**

**Chemistry**

**General considerations**

All chemical reagents are in analytical grade, obtained from commercial suppliers, and used without further purification. Reactions were monitored by thin layer chromatography (TLC) carried out on 0.25 mm silica plates (Wish Chemical, Yantai, China), using UV light as the visualizing agent and an ethanolic solution of ammonium molybdate and heat, or ninhydrin and heat as developing agents. Preparative TLC was carried out on 0.5 mm silica plates (Wish Chemical, Yantai, China). If not specially mentioned, flash column chromatography uses silica gel (200-300 mesh) supplied by Tsingtao Haiyang Chemicals (China).

Chromatographic analysis was performed using 1260 infinity II analytical HPLC system (Agilent) equipped with a Poroshell 120 EC-C18 column (2.7 μm, 3.0×150 mm). UPLC-MS analysis was performed on an ACQUITY UPLC I-Class SQD 2(Waters) system equipped with electrospray ionization (ESI) and a BEH C18 Acquity column (1.7 μm, 2.1×50 mm). High resolution mass spectra (HRMS) were recorded on a Fourier Transform Ion Cyclotron Resonance Mass Spectrometer (Solarix XR, Bruker).

NMR spectra were recorded on Bruker-500 MHz NMR (AVANCE III), Bruker-400 MHz NMR (AVANCE III) or Bruker-600 MHz (AVANCE Neo) spectrometers and evaluated using MestReNova (Mestrelab Research) software. TMS was used as internal standard for ^1^H NMR (0.00 ppm), and solvent signal was used as reference for ^1^H NMR (CDCl_3_, 7.26 ppm), ^13^C NMR (CDCl_3_, 77.16 ppm). The following abbreviations were used to explain the multiplicities: s = singlet, d = doublet, t = triplet, q = quartet, m = multiplet, br = broad.

**Synthesis of MNP-caged QM probe**

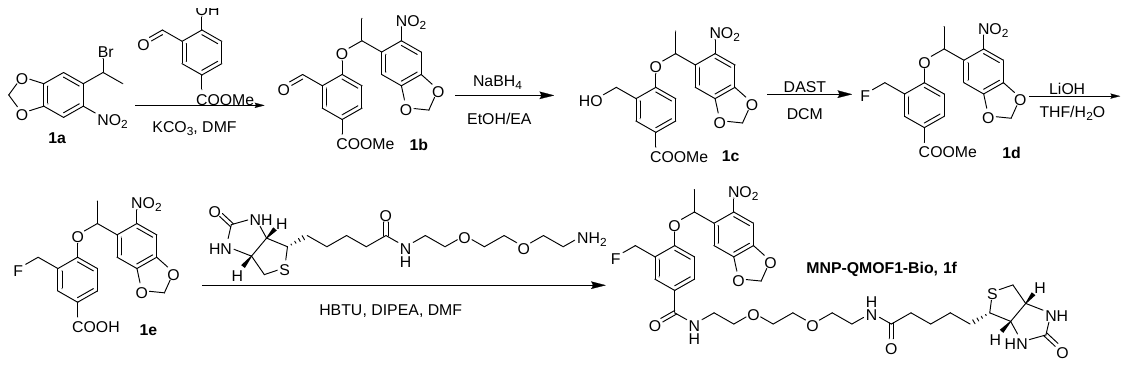

Compounds 1a^1^, 1b^2^, 1c^3^, 1d^4^, 1e^4^ and 1f^4^ were prepared according to reported procedures.

Compound 1b

To a stirring solution of methyl 3-formyl-4-hydroxybenzoate (0.9 g, 5 mmol) and 1a (1.4 g, 1.0 eq.) in DMF (20 mL) was added K_2_CO_3_ (1.38 g, 2.0 eq.). After being stirred at room temperature overnight, the mixture was quenched by saturated NH_4_Cl (80 mL) and extracted with EtOAc (30 mL, three times). The combined organic phase was dried over Na_2_SO_4_ and concentrated on a rotary evaporator. The crude mixture was purified by Flash Chromatography on SiO_2_ gel (elution with PE/EtOAc= 4/1) to afford 2b as a white solid (1.7 g, 91%). ^1^H NMR (400 MHz, Chloroform-d) δ 10.60 (s, 1H), 8.49 (d, J = 2.3 Hz, 1H), 8.08 (dd, J = 8.8, 2.3 Hz, 1H), 7.59 (s, 1H), 7.11 (s, 1H), 6.76 (d, J = 8.8 Hz, 1H), 6.30 (q, J = 6.2 Hz, 1H), 6.11 (dd, J = 14.1, 1.2 Hz, 2H), 3.88 (s, 3H), 1.80 (d, J = 6.2 Hz, 3H); ^13^C NMR (126 MHz, CDCl_3_) δ 188.44, 165.66, 162.18, 153.08, 147.69, 141.16, 136.96, 135.07, 131.01, 124.81, 123.45, 113.42, 105.64, 105.58, 103.25, 77.25, 77.00, 76.74, 72.93, 52.12, 23.12.

Compound 1c

In a round-bottomed flask, compound 1b (1.5 g, 4 mmol) was diluted in EA (30 mL), then a suspension of NaBH_4_ (95 mg, 0.66 eq.) in 6 mL of absolute EtOH was added. The reaction mixture was allowed to stir for 30 minutes at room temperature. The reaction mixture was quenched with aq. 10% NaOH and stirred until the solution was homogeneous. Water was added and ethanol was evaporated under reduced pressure. The aqueous mixture was extracted with CH_2_Cl_2_ (3x) and the combined organic extracts were washed with NaHCO_3_ (aq. 5%), then water. The solution was dried over anhydrous Na_2_SO_4_, filtered and concentrated in vacuo. The crude mixture was purified by Flash Chromatography on SiO_2_ gel (elution with PE/EtOAc= 1/1) to afford 2b as a white solid (1.4 g, 93%). ^1^H NMR (400 MHz, Chloroform-d) δ 8.03 (d, J = 2.2 Hz, 1H), 7.80 (dd, J = 8.6, 2.3 Hz, 1H), 7.57 (s, 1H), 7.07 (s, 1H), 6.57 (d, J = 8.6 Hz, 1H), 6.19 (q, J = 6.2 Hz, 1H), 6.08 (dd, J = 16.5, 1.2 Hz, 2H), 4.83 (s, 2H), 3.85 (s, 3H), 2.29 (s, 1H), 1.73 (d, J = 6.2 Hz, 3H); ^13^C NMR (126 MHz, CDCl_3_) δ 166.61, 158.28, 153.02, 147.51, 141.28, 135.95, 131.00, 130.18, 129.59, 123.02, 111.77, 105.71, 105.56, 103.16, 72.07, 61.14, 51.93, 23.35.

Compound 1d

Diethylaminosulfur trifluoride (528 µL, 2.0 eq.) was added to a solution of compound 1d (750 mg, 2 mmol) in DCM (15 mL) at 0^o^C. The reaction mixture was stirred at r.t. overnight, washed with H_2_O (20 mL ×3) and brine (20 mL ×1). The crude product was purified by column chromatography (SiO_2_, PE/DCM = 1/1) to afford 1d (656 mg, 87%). ^1^H NMR (600 MHz, Chloroform-*d*) δ 8.05 (t, *J* = 2.0 Hz, 1H), 7.89 – 7.86 (m, 1H), 7.56 (s, 1H), 7.07 (s, 1H), 6.62 (dd, *J* = 8.8, 1.2 Hz, 1H), 6.20 (q, *J* = 6.2 Hz, 1H), 6.07 (dd, *J* = 22.2, 1.3 Hz, 2H), 5.54 (ddd, *J* = 94.7, 47.6, 11.0 Hz, 2H), 3.86 (s, 3H), 1.73 (d, *J* = 6.2 Hz, 3H); ^13^C NMR (151 MHz, CDCl_3_) δ 166.33, 158.50, 158.48, 153.07, 147.54, 141.31, 135.93, 132.35, 132.34, 131.39, 131.34, 125.21, 125.09, 123.03, 111.97, 105.84, 105.51, 103.16, 80.74, 79.64, 72.11, 51.99, 23.29.

Compound 1e

1 M LiOH aqueous solution (600 µL, 0.6 mmol) was added to a solution of the compound 1d (73 mg, 0.2 mmol) in THF/H_2_O (2.4 mL/0.8 mL) at 0^o^C. The reaction mixture was stirred at r.t. overnight, diluted with 4 mL of H_2_O, neutralized with 1M HCl on ice, extracted with EtOAc, and dried over Na_2_SO_4_. The crude product was used in next step without further purification.

Compound 1f

To a solution of the carboxylic acid in dry DMF (3 mL), HBTU (83 mg, 1.1 eq.), DIEA (99 µL, 3.0 eq.), and Biotin-PEG2-NH_2_ (75 mg, 1.0 eq.) were added. The reaction mixture was stirred at r.t. under Ar overnight, the mixture was diluted with EtOAc (15 mL), and washed sequentially with deionized water (15 mL), saturated NaHCO_3_ (15 mL) and brine (15 mL). The organic phase was dried over Na_2_SO_4_ and concentrated on a rotary evaporator. The residue was purified by preparative TLC using silica plates (DCM/MeOH = 10/1) to afford 1f as white gel (86 mg, 60 %).

^1^H NMR (400 MHz, Chloroform-d) δ 7.87 (s, 1H), 7.70 (d, J = 8.6 Hz, 1H), 7.55 (s, 1H), 7.21 – 7.13 (m, 1H), 7.06 (s, 1H), 6.68 – 6.56 (m, 2H), 6.54 (s, 1H), 6.17 (q, J = 6.1 Hz, 1H), 6.09 (d, J = 7.4 Hz, 2H), 5.73 – 5.35 (m, 3H), 4.47 (dd, J = 7.9, 4.8 Hz, 1H), 4.27 (dd, J = 7.9, 4.6 Hz, 1H), 3.72 – 3.49 (m, 10H), 3.45 – 3.29 (m, 2H), 3.11 (td, J = 9.1, 7.5, 4.0 Hz, 1H), 2.87 (dd, J = 12.9, 4.8 Hz, 1H), 2.70 (d, J = 12.8 Hz, 1H), 2.16 (t, J = 7.5 Hz, 2H), 1.72 (d, J = 6.2 Hz, 3H), 1.63 (dd, J = 13.1, 5.8 Hz, 2H), 1.40 (hept, J = 7.6 Hz, 4H); ^13^C NMR (151 MHz, CDCl_3_) δ 173.53, 166.75, 164.09, 157.27, 157.25, 153.09, 147.53, 141.28, 136.05, 129.66, 129.64, 129.62, 128.92, 128.87, 127.17, 125.19, 125.08, 112.11, 105.84, 105.46, 103.23, 80.88, 79.78, 72.04, 70.11, 70.03, 69.92, 69.87, 61.80, 60.23, 55.56, 40.46, 39.85, 39.10, 35.87, 28.12, 28.00, 25.54, 23.28; ^19^F NMR (471 MHz, CDCl_3_) δ -215.22; **FT-HRMS (ESI)** m/z calcd. for C_33_H_43_FN_5_O_10_S [M+H]^+^: 720.2709; Found: 720.2709.

**Synthesis of PAB-caged QM probe**

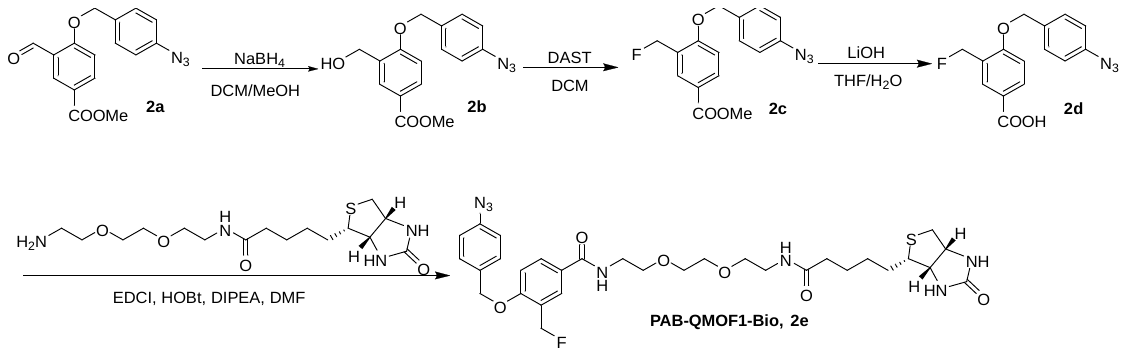

Compounds 2a^2^, 2b^5^, 2c^4^, 2d^4^ and 2e^2^ were prepared according to reported procedures.

Compound 2b

To a solution of compound 2b (940 mg, 3 mmol) in 2 mL of MeOH and 20 mL of DCM at 0 °C was added NaBH_4_ (114 mg, 1.0 eq.). The resulting mixture was then stirred at 0 °C for 15 minutes then 30 additional minutes at room temperature under argon. After the reaction was completed, the reaction was stopped by adding 30 mL of a saturated solution of NaHCO_3_ and the phases were separated. The aqueous phase was extracted twice with 30 mL of CH_2_Cl_2_, the organic layers were combined, dry over MgSO_4_ and evaporated to dryness. The crude mixture was purified by Flash Chromatography on SiO_2_ gel (elution with PE/EtOAc= 1/1) to afford 2b as a white solid (900 mg, 95%). ^1^H NMR (400 MHz, Chloroform-*d*) δ 8.00 – 7.86 (m, 2H), 7.33 (d, *J* = 8.5 Hz, 2H), 6.99 (d, *J* = 8.5 Hz, 2H), 6.87 (d, *J* = 8.6 Hz, 1H), 5.06 (s, 2H), 4.67 (s, 2H), 3.81 (s, 3H), 2.12 (s, 1H); ^13^C NMR (151 MHz, CDCl_3_) δ 165.69, 158.84, 139.22, 131.65, 130.09, 129.12, 128.52, 127.95, 121.96, 118.38, 110.01, 68.72, 60.34, 50.92.

Compound 2c

Diethylaminosulfur trifluoride (544 µL, 2.0 eq.) was added to a solution of the compound 2b (630 mg, 2.0 mmol) in DCM (15 mL) at 0 ^o^C. The reaction mixture was stirred at r.t. overnight, washed with H_2_O (20 mL ×3) and brine (20 mL ×1). The crude product was purified by column chromatography (SiO_2_, Hexane/EtOAc= 2/1) to afford 2c (420 mg, 84%). 1H NMR (400 MHz, Chloroform-d) δ 8.06 – 7.91 (m, 2H), 7.33 (d, J = 8.5 Hz, 2H), 6.98 (d, J = 8.5 Hz, 2H), 6.88 (d, J = 9.9 Hz, 1H), 5.47 (s, 1H), 5.35 (s, 1H), 5.06 (s, 2H), 3.82 (s, 3H); ^13^C NMR (151 MHz, CDCl3) δ 165.48, 158.66, 158.64, 139.11, 131.66, 131.18, 131.16, 129.81, 129.76, 127.82, 124.26, 124.15, 121.92, 118.31, 110.20, 79.49, 78.39, 76.21, 76.00, 75.79, 68.71, 50.97.

Compound 2d

1 M LiOH aqueous solution (600 µL, 0.6 mmol) was added to a solution of the 2c (0.2 mmol) in THF/H_2_O (2.4 mL/0.8 mL) at 0^o^C. The reaction mixture was stirred at r.t. overnight, diluted with 4 mL of H_2_O, neutralized with 1M HCl on ice, extracted with EtOAc, and dried over Na_2_SO_4_. The crude product was used in next step without further purification.

Compound 2e

To a solution of the carboxylic acid in dry DMF (3 mL), EDCI (38.2 mg, 1.0 eq.) and HOBt (27 mg, 1.0 eq), and DIEA (99 µL, 3.0 eq.) were added. The resulting mixture was stirred at 0 ℃ under Ar for 1 h, followed by the addition of Biotin-PEG2-NH_2_ (1.0 eq.). After being stirred at room temperature overnight, the mixture was diluted with EtOAc (15 mL), and washed sequentially with deionized water (15 mL), saturated NaHCO_3_ (15 mL) and brine (15 mL). The organic phase was dried over Na_2_SO_4_ and concentrated on a rotary evaporator. The residue was purified by preparative TLC using silica plates (DCM/MeOH = 10/1) to afford 2e as white gel (94 mg, 71 %). ^1^H NMR (400 MHz, Chloroform-*d*) δ 7.85 (d, *J* = 10.9 Hz, 2H), 7.39 (d, *J* = 8.5 Hz, 2H), 7.19 (s, 1H), 7.04 (d, *J* = 8.4 Hz, 2H), 6.95 (d, *J* = 8.4 Hz, 1H), 6.64 (s, 1H), 6.33 (s, 1H), 5.53 (s, 1H), 5.41 (s, 1H), 5.10 (s, 2H), 4.42 (s, 1H), 4.23 (s, 1H), 3.71 – 3.31 (m, 12H), 3.12 – 3.02 (m, 1H), 2.84 (dd, *J* = 12.9, 4.8 Hz, 1H), 2.66 (d, *J* = 12.8 Hz, 1H), 2.14 (t, *J* = 7.5 Hz, 2H), 1.59 (dq, *J* = 14.6, 7.3, 6.9 Hz, 4H), 1.35 (t, *J* = 7.8 Hz, 2H); ^13^C NMR (151 MHz, CDCl_3_) δ 173.65, 167.05, 164.03, 158.52, 158.49, 140.06, 132.87, 129.70, 129.68, 128.89, 128.05, 128.00, 126.91, 125.19, 125.08, 119.31, 111.52, 80.70, 79.60, 70.08, 70.04, 70.01, 69.71, 61.80, 60.25, 55.37, 40.45, 39.84, 39.16, 39.14, 35.77, 27.99, 27.96, 25.42; ^19^F NMR (471 MHz, Chloroform-d) δ -213.32 (d, J = 10.6 Hz); **FT-HRMS (ESI)** m/z calcd. for C_31_H_41_FN_7_O_6_S [M+H]^+^: 658.2819; Found: 658.2822.

**Synthesis of photocatalysts (Ir_3_-PEG_4_-NHS)**

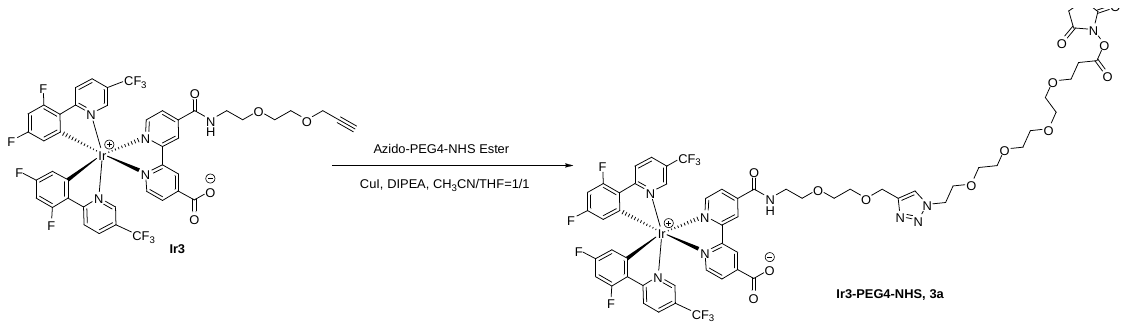

Compounds Ir3^6^ and Ir_3_-PEG_4_-NHS^7^ were prepared according to reported procedures.

In an oven-dry 25 mL round-bottom flask, Ir3 (30 mg, 0.028 mmol), Azido-PEG4-NHS Ester (11 g, 1.0 eq.) and copper(I) iodide (5.3 mg, 1.0 eq.) were dissolved in a 50:50 mixture of THF and MeCN (4 mL). DIPEA (4.6 μL, 1.0 eq.) was added and the reaction mixture was vigorously stirred at room temperature for 3 h under an argon atmosphere. Ir_3_-PEG_4_-NHS (30 mg, 73 %) was obtained after purification by HPLC (A/B = 5/95 to 95/5 over 35 min, A = CH_3_CN, B = H_2_O). ^1^H NMR (400 MHz, Chloroform-*d*) δ 8.47 (d, *J* = 9.9 Hz, 3H), 8.12 (dd, *J* = 12.6, 5.7 Hz, 2H), 8.07 – 7.94 (m, 6H), 7.89 (s, 2H), 7.71 (s, 2H), 7.64 (s, 2H), 6.71 – 6.62 (m, 3H), 5.65 (dd, *J* = 14.3, 8.0 Hz, 3H), 3.84 (q, *J* = 6.5, 5.2 Hz, 4H), 3.65 – 3.59 (m, 18H), 2.88 (t, *J* = 6.4 Hz, 2H), 2.83 (s, 4H); ^13^C NMR (151 MHz, CDCl_3_) δ 169.04, 167.79, 167.75, 166.76, 165.68, 164.03, 163.95, 163.92, 163.48, 163.40, 163.34, 163.04, 161.74, 161.65, 156.97, 154.88, 154.60, 150.22, 149.97, 146.47, 145.83, 145.59, 145.17, 136.59, 127.90, 126.39, 126.27, 126.16, 124.60, 124.17, 123.73, 123.64, 123.50, 122.38, 122.36, 120.57, 120.55, 114.26, 114.15, 114.09, 113.97, 100.26, 100.18, 100.09, 99.91, 70.81, 70.70, 70.60, 70.54, 70.51, 70.46, 70.43, 70.34, 70.19, 69.59, 69.36, 69.27, 69.20, 65.71, 40.09, 32.14, 25.60; **FT-HRMS (ESI)** m/z calcd. for C_58_H_52_F_10_IrN_9_O_13_ [M+H]^+^: 1466.3227; Found: 1466.3221.

**Synthesis of nanobody-photocatalyst conjugate (ZHER-Ir_3_)**

ZHER-[Ir]_3_ conjugate was prepared in two steps.

In a 1.5-mL centrifuge tube, 760.4 μL of water was added firstly. Then 200 μL of 6.7 mg/mL ZHER-GGS-Flag-GGS-LPETG-His in PBS (pH 7.4), 518 μL of a 2 mg/mL solution of GGG-[N_3_]_3_, 120 μL of 108 μM mgSrtA, and 1.6 μL of 1M CaCl_2_ solution were added in mixture orderly. After immediate vortexing, the resulting mixture was incubated at room temperature for 30 min. Then the mixture was quenched by adding 16 μL 200 mM MESET solution and incubated for 10 min. The crude sample was desalted using AKTA according to general protocol. After that, crude product sample was divided into several tubes and 48 μL His smart beads were added per milliliter. The mixture was rotated on sky wheel for 20 min and centrifuged (13,000 g) for 5 min, then the supernatant was all collected for final ultrafiltration and concentration using 3 kDa Amicon Ultra-centrifugal Filter (Millipore). The product ZHER-[N_3_]_3_ could be detected by LC-MS.

The preparation of ZHER-[Ir]_3_ from ZHER-[N_3_]_3_ was performed using Click-It kit (Thermo, C10276) according to the manufacture’s protocol with minor modifications. In brief, 47.7 μL of 1 mg/mL ZHER-[N_3_]_3_ was treated with 100 μL reaction buffer A and 3.33 μL of a 30 mM solution of Ir-azide in DMSO. Then the mixture was diluted with ddH_2_O to 160 μL, followed by the addition of 10 μL of 40 mM CuSO_4_ (component B) and 10 μL of component C. The mixture was vortexed and incubated for 4 min at room temperature, followed by the addition of 18 μL of component D. The resulting mixture was then incubated at 30ºC for 25 min in a ThermoMixer, protected from light. The reaction mixture was passed through DPBS-equilibrated Micro Bio-Spin P-6 columns (Bio-Rad) (100 μL reaction mixture per column) to yield ZHER-[Ir]_3_ in DPBS. The number-average degree of labeling (DOL) for photocatalyst moiety determined by LC-MS was normally ~3.

**HPLC analysis of photocatalytic decaging of PAB-QM-Bio**

PAB-QM-Bio (0.1 mM) was mixed with butylamine (100 mM), NADH (0.5 mM) and Ir8 (10 mol%) in 10:1 DMSO/PBS (pH 7.4) buffer, and irradiated by mild blue LED (4 mW/cm^2^) for 0 min, 2 min, 5 min, 10 min and 20 min at room temperature. Groups without light irradiation or Ir8 were used as controls. Reactions were analyzed on an ACQUITY UPLC I-Class (Waters) system equipped with a BEH C18 Acquity column (1.7 μm, 2.1×50 mm). Peaks were detected using 293-nm detector. Mobile phases: A (MeCN), B (5% MeCN in H_2_O). Flow rate = 0.3 mL/min; Eluent A/B gradually changed from 5/95 to 100/0 in 4 min.

**Photocatalytic decaging of PAB-QM-Bio for BSA labeling**

BSA (3 mg/mL) was dissolved in PBS (pH 7.4). Ir (5.0 μM) and NADH (1500 μM) were added in mixtures and vortexed. Then the mixtures were divided in two parts and combined with PAB-OF1-QM-Bio probe (100 μM) and PAB-OF2-QM-Bio probe (100 μM) respectively to afford reaction mixtures with total solution volume of 1500 μL each. The final solution was added in a 96-well plate 230 μL per well. The samples were then irradiated by mild blue LED (4 mW/cm^2^) for different time scales (0, 1, 3, 5, 10, 15min) at room temperature. After that, 200 μL of each sample was taken and combined with 400 μL methanol and 100 μL chloroform overnight in -30ºC. Each precipitate protein sample was washed with methanol three times and finally dissolved in 300μL 1.2% SDS/PBS buffer. 10 μL of each sample was taken and combined with 30 μL water and 10 μL reducing loading buffer (CWBIO) followed by heating at 95°C for 10 minutes. Immunoblotting was performed with 4-15% gradient SDS-PAGE and 0.2 μm PVDF membrane according to the general protocol. Mouse anti-biotin mAb (Santa Cruz, sc-101339, 1:1,000 dilution) and HRP-linked anti-mouse IgG (Cell Signaling Technology, 7076S, 1:5,000 dilution) were used as primary and secondary antibodies, respectively. The taken aliquots were also analyzed by Coomassie blue staining for protein quantification.

**Biology**

**General considerations**

Media, buffer components, kits, and cloning enzymes were used as received from the specified commercial suppliers. All expression media, buffers and antibiotics were prepared using purified H_2_O (Mili-Q Reference) and autoclaved or filter-sterilized, as appropriate. All the flow cytometry antibodies were purchased from biolegend.

Images of protein gels including Coomassie SDS-PAGE gel and immunoblotting membranes were taken on ChemiDoc XRS+ (Bio-Rad). Fluorescence imaging of SDS-PAGE gel was performed on Typhoon FLA 9500 (GE healthcare). Flowcytometry was performed on LSRFortessa™ Cell Analyzer (BD biosciences) supplemented with BD FACSDiva software.

Statistical analyses were performed using GraphPad Prism software 8.0.2. Image Lab was used for

western blotting analysis. Flow cytometric analysis was performed using FlowJo version 10.6.2. Comparisons over groups were analyzed by two-way ANOVA tests followed by Sidak's multiple comparisons test and two-tailed unpaired t test. In all figures, ns, p > 0.05; * p < 0.05; ** p < 0.01; *** p < 0.001; **** p < 0.0001

The experimental schemes were created with BioRender.com (agreement number OF257OX2EO).

**Cell line culture and Mice**

HEK293T, HLA-A2:02*01 K562, 1G4-JC5, MC38 and MC38 B2M KO cells were maintained in DMEM (Gibco) supplemented with 10% FBS (v/v., BI), 100 IU/ml penicillin and 100 mg/ml streptomycin (Thermo Fisher). All of them were cultured in an incubator at 37°C under 5% CO2. C57BL/6J mice were purchased from purchased from Charles River. OT-I+/+ mice with a C57BL/6J genetic background were gifts from Meng Xv’s lab. Both male and female mice of 8-15 weeks of age were used for most experiments. All purchased mice were specific pathogen-free (SPF) animals and were bred or housed under clean conditions. All animal procedures were performed in accordance with the Guidelines for Care and Use of Laboratory Animals published in GB/T 35892-2018 and the experiments were approved by Institutional Animal Use and Care Committee of Peking University of China (CCME-ChenP-3).

**Primary cells preparation and culture**

Splenocytes were obtained from the spleen of C57BL/6 mice (purchased from Charles River) after red blood cells lysis (CWBIO) for use immediately or cryopreservation in CELLSAVING (NCM biotech) at -80°C for future use. Splenocytes were maintained in the complete T cell medium (RPMI 1640 (Gibco) with 10% FBS, 50 μM 2-Mercaptoethanol (Gibco), 10 mM HEPES (Gibco), 1 mM sodium pyruvate (Gibco) and 1$\times$Minimum Essential Medium Non-Essential Amino Acids (Gibco), 100 IU/mL penicillin and 100 mg/mL streptomycin (Gibco)) with 50 IU/mL rhIL2 (Procell).

**Photocatalytic proximity labeling of PAB-QM probes in CD40L-CD40 system**

HEK293T cells were transfected with CD40L-Tdtomato, CD40-GFP and CD45-GFP plasmids respectively for NHS-Ir based CAT-Cell, and CD40L-Tdtomato was replaced with CD40L-T2A-HER2-Tdtomato for ZHER-Ir_3_ based CAT-Cell experiment. HEK293T cells were first collected and counted. CD40L+ cells were centrifuged for 2 minutes at 450 g. The cell pellets were resuspended at a density of 5$\times$10^7^ cells per mL with 1$\times$PBS (Solarbio) containing 100 µM NHS-[Ir] and incubated in a water bath at 37°C for 5 minutes, which was quenched by cell culture medium with 10% Fetal Bovine Serum (Gibco) and centrifuged for 2 minutes at 450 g. Finally, CD40L+ cells were resuspended with DMDM (Gibco) with 1% FBS at a density of 3$\times$10^5^ cells per 75 µL with 50 µM OF1 or OF2 probe. At the same time, CD40/45+ cells were harvested, counted and resuspended with DMDM with 1% FBS at a density of 1$\times$10^5^ cells per 75 µL with 50 µM OF1 or OF2 probe.

Co-incubations were set up in 96-well V-bottom plates at a ratio of 3:1 CD40L+ cells to CD40/45+ cells (4$\times$10^5^ cells per well) and co-incubated for 60 minutes at 37°C. The co-incubation was then placed at 4°C for 10 minutes and NADH (GPC Bio) was added to the culture medium with the final concentration of 7 mg/mL followed another incubation at 4°C for 5 minutes. After the cell medium cooling down, it was irradiated by direct 465 nm light for 5 minutes on the ice and then rest in the dark at 4°C for 15 minutes. The cell culture was then centrifuged for 2 minutes at 450 g. After washing four times with FACS buffer (1$\times$PBS with 2% FBS), cells were stained with streptavidin-APC at room temperature for 30 minutes. After washing twice, cell mixtures were analyzed by BD LSRFortessa™ Cell Analyzer (BD Bioscience).

**Photocatalytic proximity labeling in pMHC-TCR system**

Lyophilized peptides were reconstituted in dimethyl sulfoxide to 10 mg/mL. For the proximity labeling assay, K562-HLA A2 cells were first counted and pulsed with 1 µg/µL each of the epitope peptides in an Eppendorf tube at 37℃ for 1 hour. After incubation, K562-HLA A2 cells were centrifuged for 2 minutes at 450 g. The cell pellets were resuspended at a density of 5$\times$10^7^ cells per mL with 1$\times$PBS (Solarbio) containing 100 µM NHS-[Ir] and incubated in a water bath at 37℃ for 5 minutes, which was quenched by cell culture medium with 10% Fetal Bovine Serum (Gibco) and centrifuged for 2 minutes at 450 g. Finally, K562-HLA A2 cells were resuspended with IMDM (Gibco) with 1% FBS at a density of 3$\times$10^5^ cells per 75 µL with 50 µM OF1. At the same time, 1G4-JC5 cells were harvested, counted and resuspended with IMDM with 1% FBS at a density of 1$\times$10^5^ cells per 75 µL with 50 µM OF1.

For experiments using 1G4-JC5 and K562-HLA A2, co-incubations were set up in 96-well V-bottom plates at a ratio of 3:1 K562 to JC5 (4$\times$10^5^ cells per well) and co-incubated for 90 minutes at 37°C. The co-incubation was then placed at 4℃ for 10 minutes and NADH (GPC Bio) was added to the culture medium with the final concentration of 7 mg/mL followed another incubation at 4℃ for 5 minutes. After the cell medium cooling down, it was irradiated by direct 465 nm light for 5 minutes on the ice and then rest in the dark at 4℃ for 15 minutes. The cell culture was then centrifuged for 2 minutes at 450 g. After washing four times with FACS buffer (1$\times$PBS with 2% FBS), cells were stained with streptavidin-APC at room temperature for 45 minutes. After washing twice, cell mixtures were analyzed by BD LSRFortessa™ Cell Analyzer (BD Bioscience). Gating strategy was as follows: cells (SSC-A versus FSC-A), single cells (FSC-W versus FSC-A), 1G4-JC5 (SSC-A versus tdTomato-PE), labelled JC5 cells (tdTomato-PE versus streptavidin-APC), activated JC5 cells (tdTomato-PE versus GFP-FITC).

For the validation of CAT-cell, prey cells were always kept at a density of 1$\times$10^5^ per well. For the validation experiments, the same protocol was used, but the relevant parameters changed accordingly.

**CAT-Cell protocol for OT-I antigen-specific T and NK cell capture**

For the co-incubation of OT-I splenocytes and MC38 cells, 3$\times$10^5^ MC38 cells were co-incubated with 1$\times$10^5^ splenocytes in a well of V-bottom 96-well plates for 2 hours at 37℃. The labeling experiment was conducted as described before. The cells were first stained with mouse Fc-blocker and Zombie Aqua^TM^ for 10 minutes at room temperature and then incubated with anti-mouse CD8a-BV421, anti-mouse CD69-PE/Cy7 and streptavidin-APC on ice for 30 min. After washing twice, cell mixtures were prepared for flow cytometry analysis.

For analysis of the specificity of CAT-cell in primary cell culture, splenocytes of C57BL/6 mice (B6 mice) were isolated and then stained with 2.5 µM CFSE (BD Bioscience) in RPMI 1640 with 1% FBS at 37℃ for 4 minutes. After that, complete cell culture medium containing 10% FBS was added to quench it. For experiment using OT-1 splenocytes and B6 splenocytes, 3$\times$10^5^ [Ir] modified peptide-pulsed MC38 cells were plated with 1$\times$10^5^ mixed splenocytes at a cell ratio of OT-1: B6 splenocytes to 1:1. The same labeling protocol was performed. The cells were stained as described and then analyzed by FACS. Gating strategy was as follows: cells (SSC-A versus FSC-A), single cells (FSC-W versus FSC-A), viable CD8+ T cells (CD8a-BV421 versus Zombie), OT-1 T cells (CD8a-BV421 versus CFSE-FITC, if needed), labelled T cells (CD8a-BV421 versus streptavidin-APC), activated CD8+ T cells (CD8a-BV421 versus CD69-PE/Cy7).

**CAT-Cell protocol for detection of antigen-specific T cells in tumor samples**

MC38-OVA and MC38-B2M KO cells were cultured in DMEM medium (Gibco), supplemented with 10% FBS, 100 IU/mL penicillin and 100 mg/mL streptomycin. C57BL/6N mice (7-8 weeks) were injected subcutaneously with 1×10^6^ tumor cells. Mice were sacrificed if the tumor diameter reached ~1 cm. The general tumor digestion protocol was referenced to a paper published by us^6^. In general, the solid tumor tissue was harvested, cut into small pieces (~2 nm) and transferred to 5 mL RPMI 1640 with 0.2 mg/mL DNAse I (Roche 10104159001), 200 UI/mL collagenase I (Worthington LS004196) and collagenase IV (Worthington LS004186). The digestion mixture was incubated in a shaker for 1 hour at 37℃. When the digestion was completed, adding 500 µL FBS to the mixture to terminate the reaction followed by passing through a 70-μm cell drainer. The single cell suspension was centrifuged at 450 g for 5 minutes and treated with 500 µL RBC lysis buffer to deplete red blood cells. Eventually, these cells were used for downstream labeling immediately. Meanwhile, cell lines MC38 and MC38-B2M KO cells were trypsin-digested for cell counting.

The labeling procedure was the same as the steps in mouse specific primary T cells capture by [Ir] functionalized tumor cells. MC38 cells were pulsed with 1 µM OVA_257-264_ peptide or GP_33-41_ peptide for 1 hour at 37℃ for detection of antigen-specific CD8+ T cells. In each reaction system, bait cells were 3×10^5^, and the prey cells (tumor single cell suspension) were 1×10^5^. After the reaction was completed, the cells were stained for cytometric analysis. The gating strategy was as follows: cells (SSC-A versus FSC-A), single cells (FSC-W versus FSC-A), viable tumor-infiltrating lymphocytes (CD45-PE versus Zombie), CD8 T cells (CD3-FITC versus CD8a-PB), NK cells (CD3-FITC versus NK1.1-PB), labeled CD8+ T cells (CD8a-PB versus streptavidin-APC), labeled NK cells (NK1.1-PB versus streptavidin-APC). And FITC anti-human PD-1, PE anti-human TIM-3, PE-Cy7 anti-human CD39, PE anti-human CD137 were used separately to characterize molecular phenotypes of Biotin+ and Biotin− CD8+ T cells.

**Supplementary Figures**

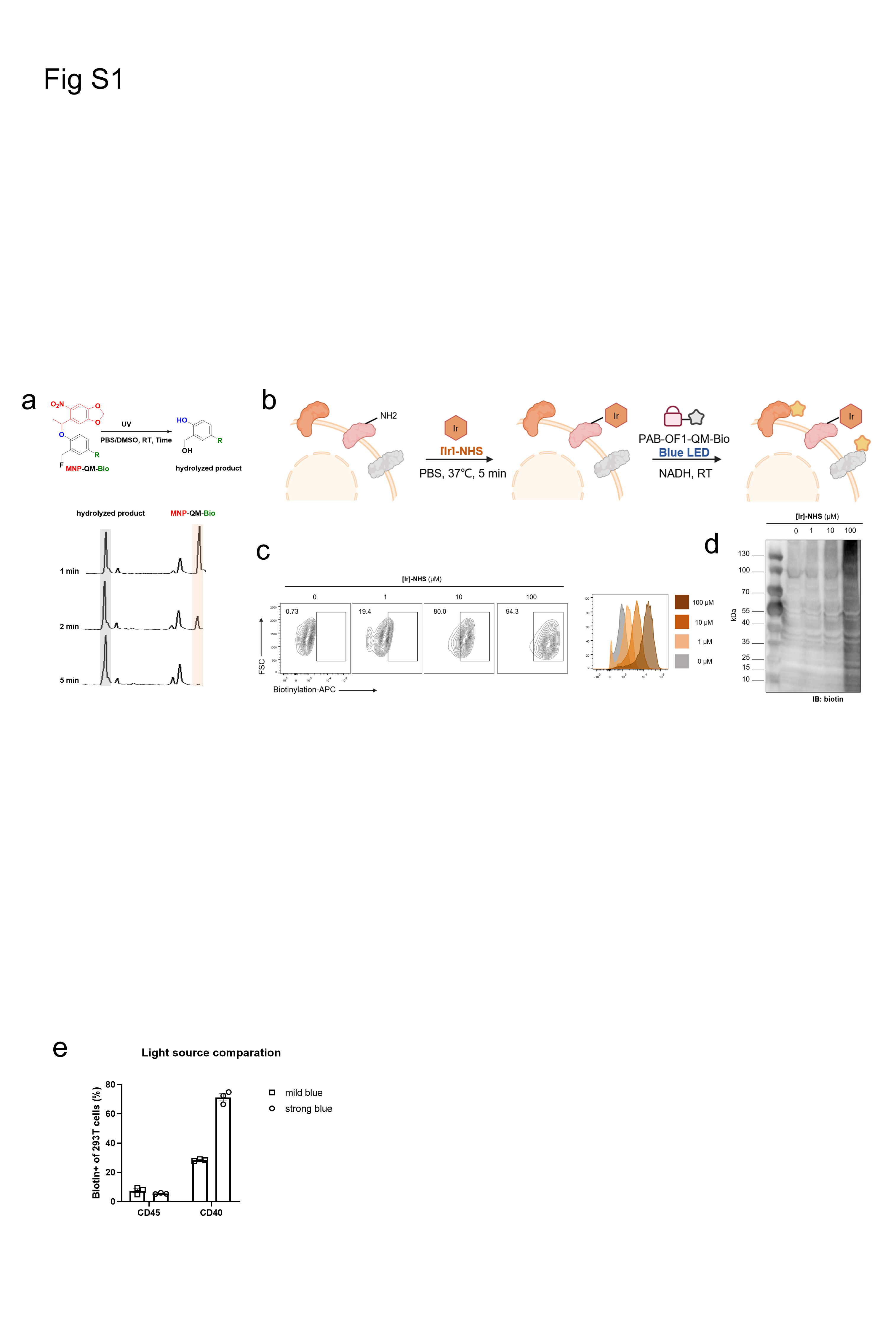

**Figure S1. NHS ester-mediated photocatalyst cell-surface functionalization.**

(a) HPLC traces of the photocatalytic decaging of MNP-QM-Bio (MNP-OF1) probe under blue LED at given time points. (b) Schematic illustration of Ir-based cis-cell labeling. (c, d) Flow cytometric analysis and western blot showing different biotinylation efficiency of different NHS-Ir treatment.

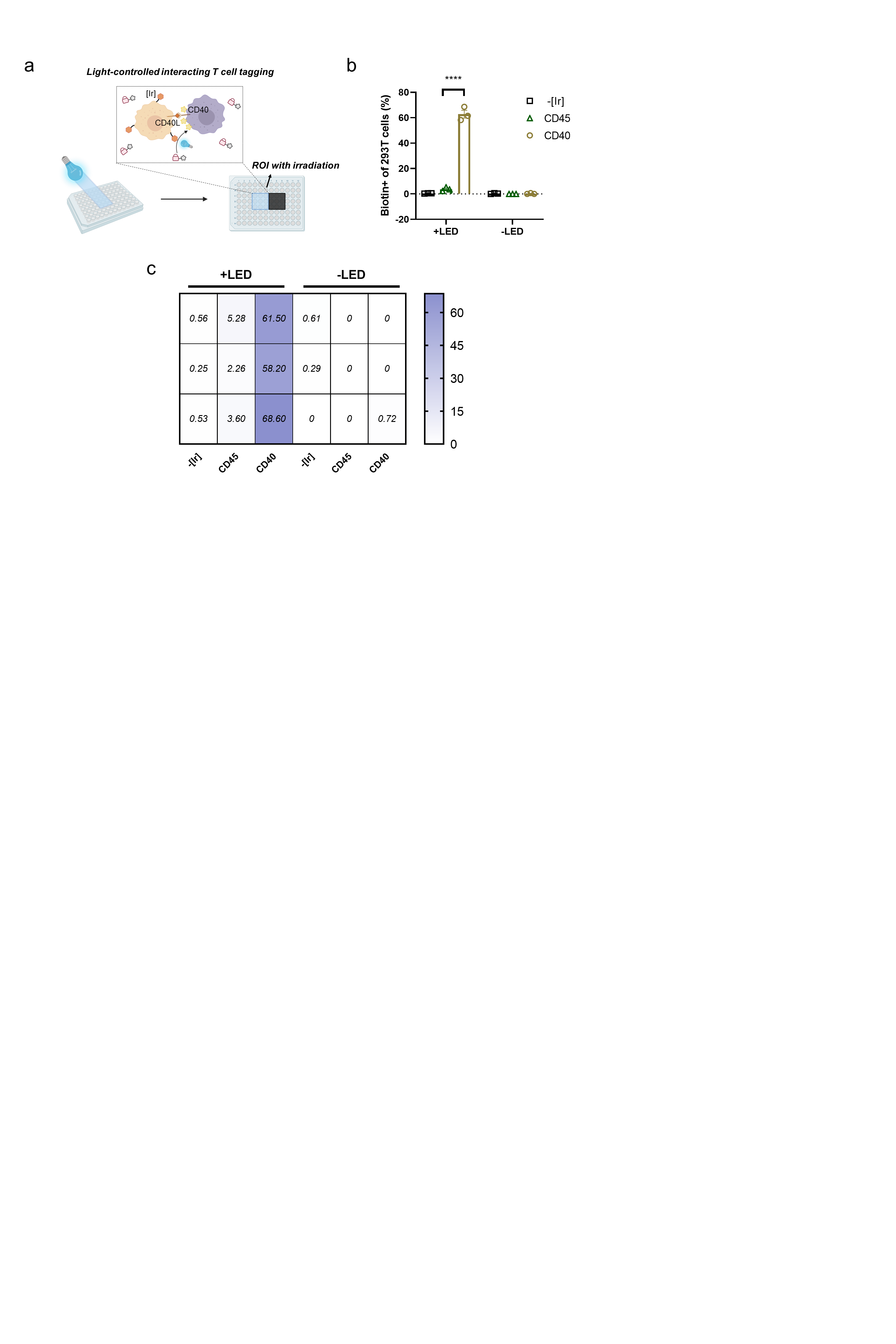

**Figure S2. CAT-Cell in combination with spatial resolution.**

(a) Schematic illustration of CAT-Cell in combination with spatial resolution. Only interacting cells which were transfected with CD40 in ROI can be biotinylation via CAT-Cell after irradiation. (b, c) Flow cytometric analysis of biotinylation carried by CAT-Cell in combination with spatial resolution. Heatmaps showing the biotinylation percentage of HEK293T cells with (left) or without (right) blue light irradiation from co-cultures in each well. The tagging percentages were calculated based on FC.

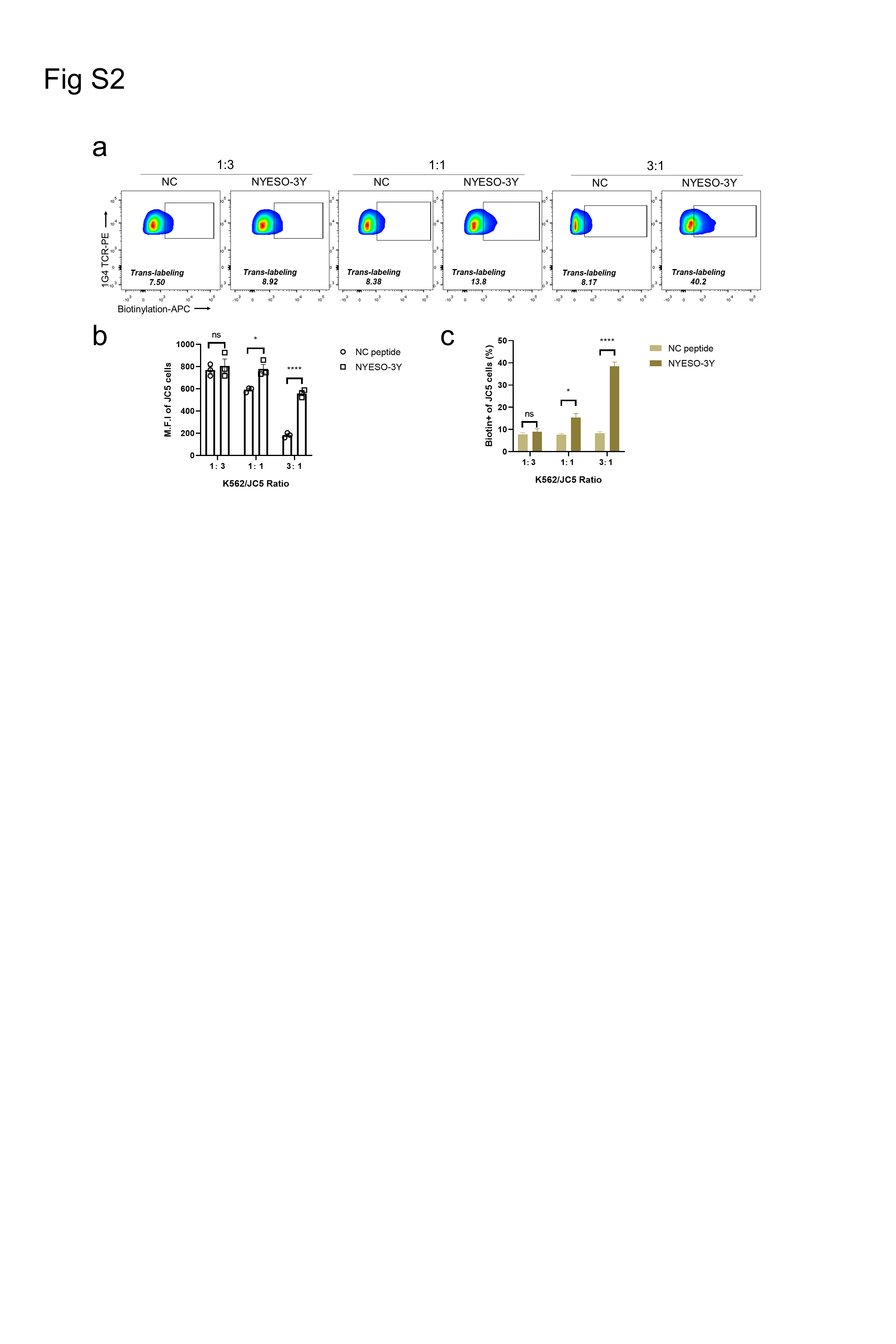

**Figure S3. Optimization of CAT-Cell applied for proximity labeling of antigen-specific T cells.**

(a) Flow cytometric plots of antigen-specific biotinylation of 1G4 TCR T cells at different ratios. (b, c) Flow cytometric analysis of biotinylation M.F.I and percentage of 1G4 TCR T cells at different ratios.

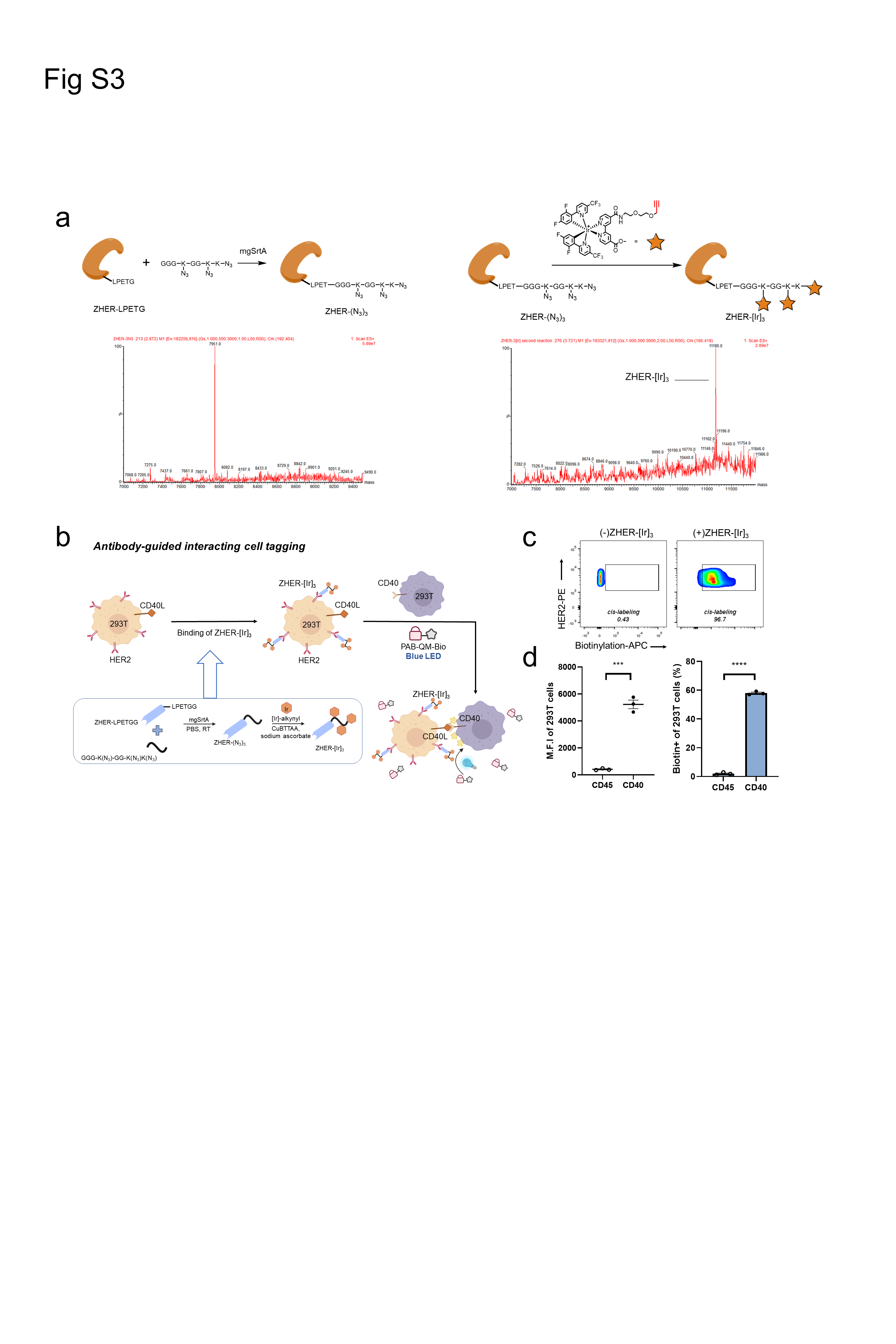

**Figure S4. Intercellular labeling via CAT-Cell in combination with nanobody-photocatalyst conjugate (ZHER-[Ir]_3_).**

(a) Synthesis and characterization of ZHER-[Ir]_3_. (b) Schematic view of ZHER-[Ir]_3_-based CAT-Cell for CD40L-CD40 interaction profiling. (c) Flow cytometric analysis showing biotinylation of cis-labeling introduced by specific binding of ZHER-[Ir]_3_ conjugate. (d) Flow cytometric analysis showing specific biotinylation of CD40-positive cells.

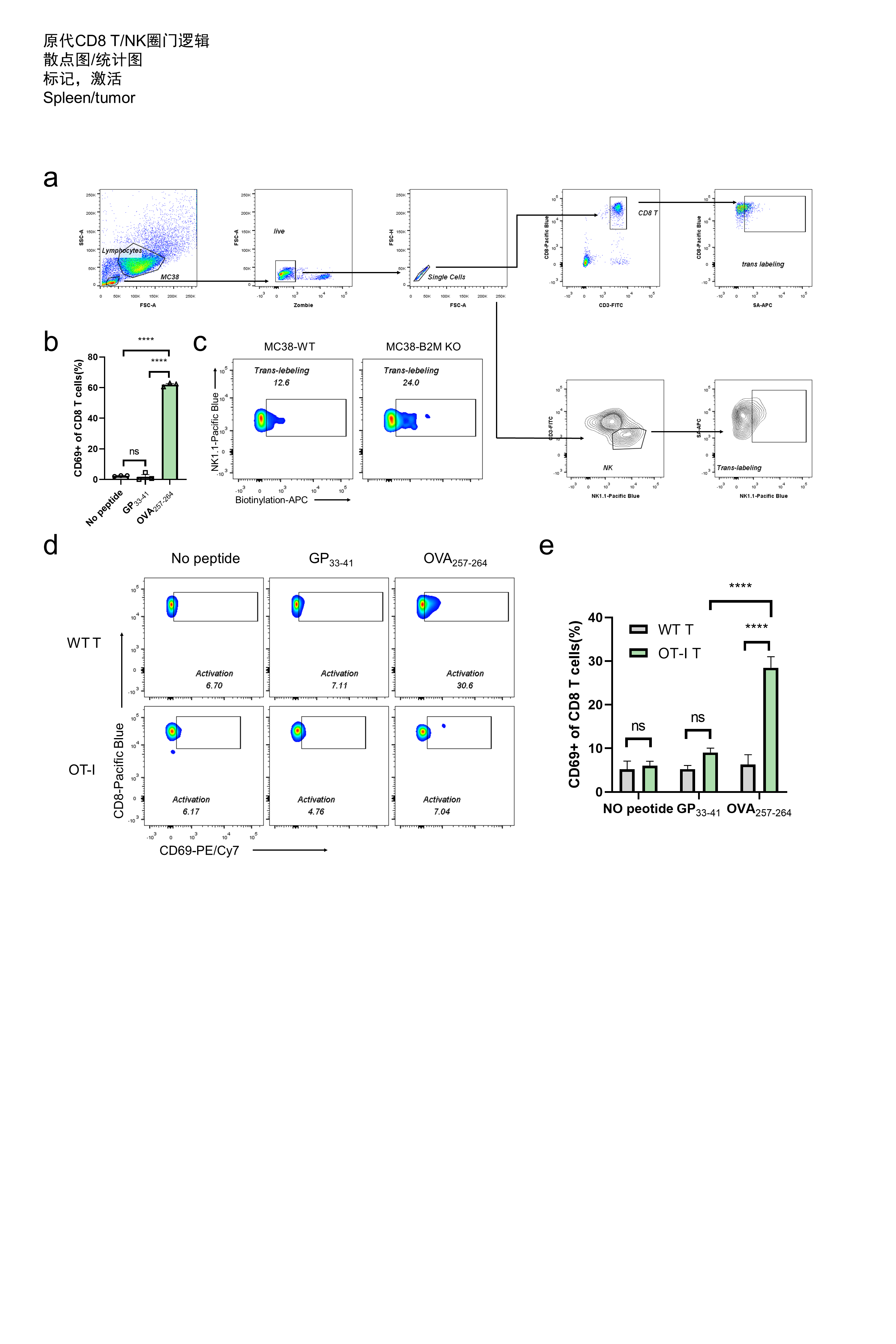

**Figure S5. CAT-Cell enabled *ex vivo* identification of antigen-specific T cells and NK cells in mouse model.**

(a) Representative flow cytometric plots showing the gating strategy of biotinylated OT-I CD8+ T cells and NK cells. (b) Flow cytometric analysis of activation degree of OT-I T. (c) Flow cytometric plots of biotinylated NK cells. (d, e) Flow cytometric analysis showing CD69 expression level of WT T and OT- I T cells in different groups in splenocyte mixture. There was only insignificant activation background under all controlled conditions, while a significant CD69 signal was detected on OT-Ⅰ T cells co-cultured with OVA_257-264_-primed MC38 cells.

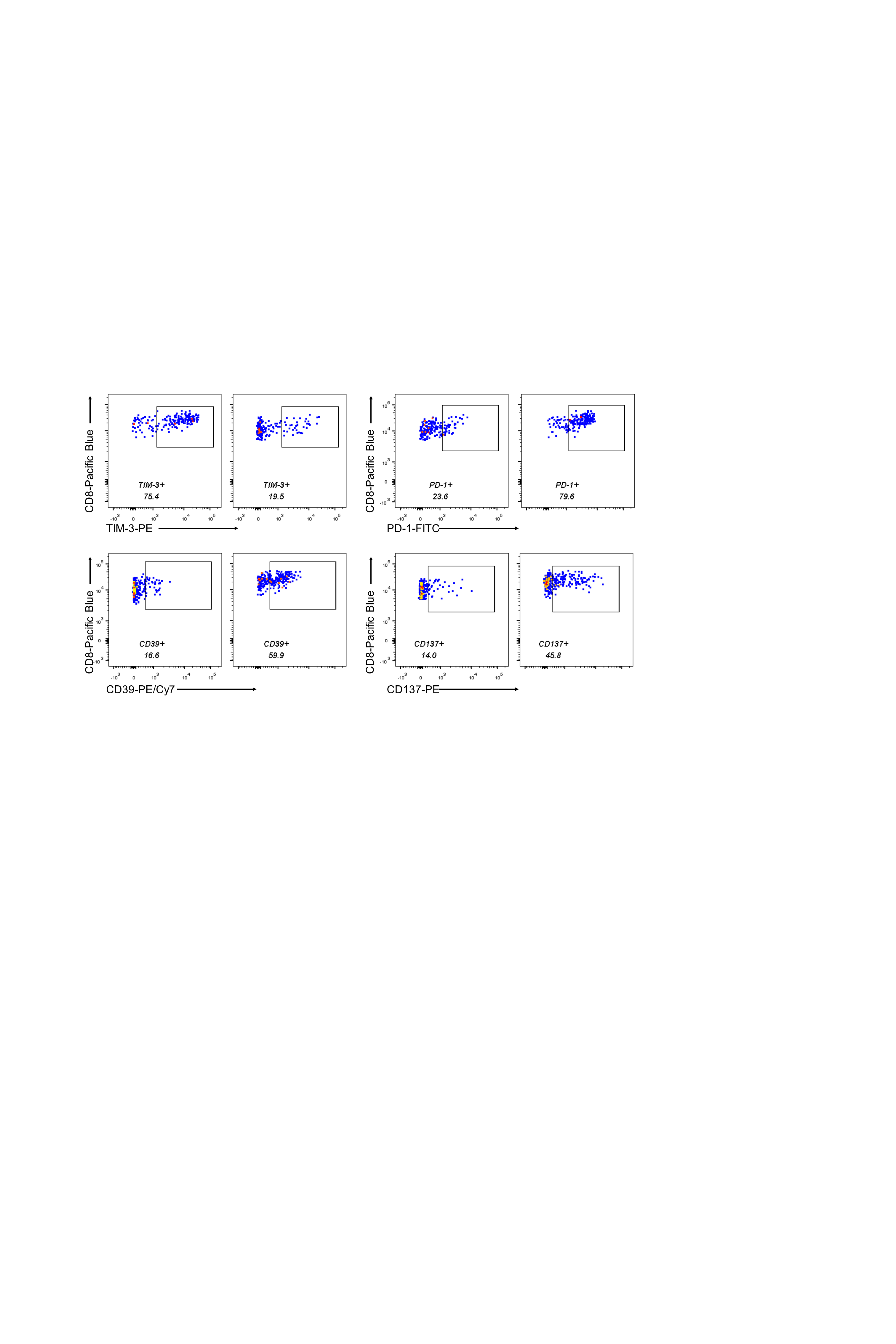

**Figure S6. Characterization of molecular phenotypes of Biotin+ and Biotin− CD8+ T cells in TILs.**

Flow cytometric plots showing the expression level of PD-1, TIM-3, CD39 and CD137 on Biotin+ CD8+ T cells and Biotin- CD8+ T cells.

**NMR SPECTRA**

**1b:** ^1^H NMR (400 MHz, Chloroform-d)

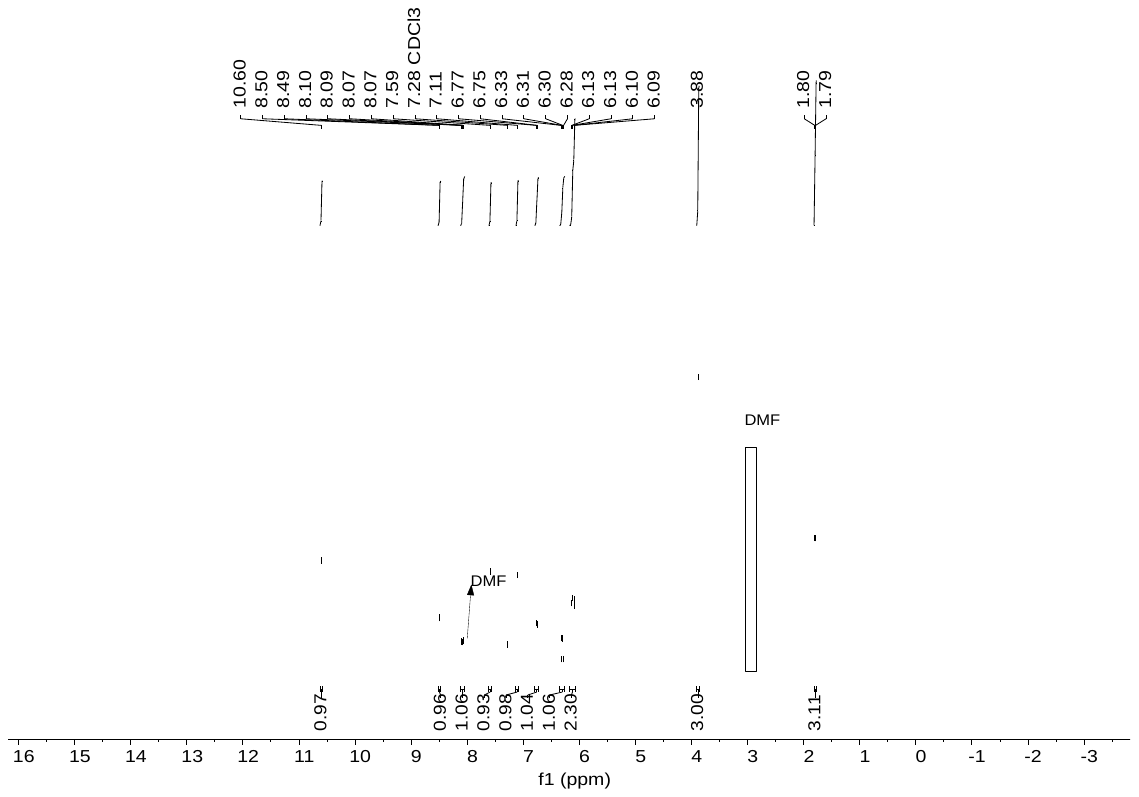

**1b:** ^13^C NMR (151 MHz, Chloroform-*d*)

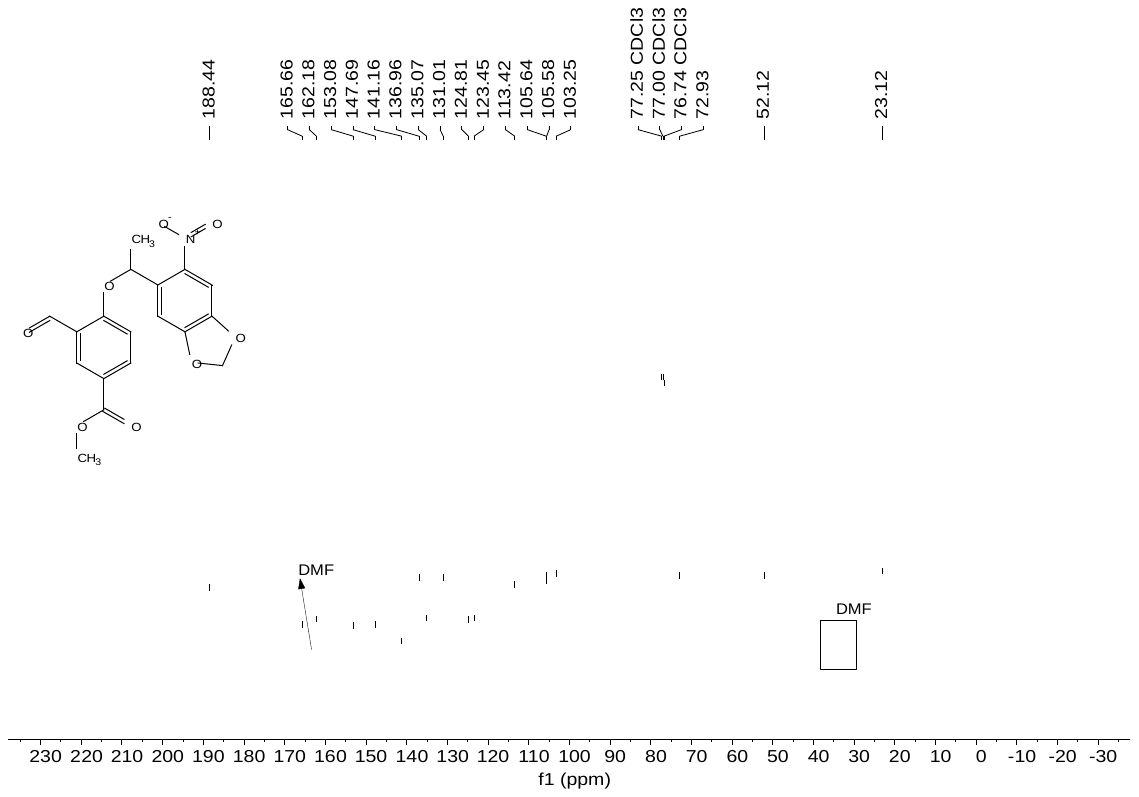

**1c:** ^1^H NMR (400 MHz, Chloroform-d)

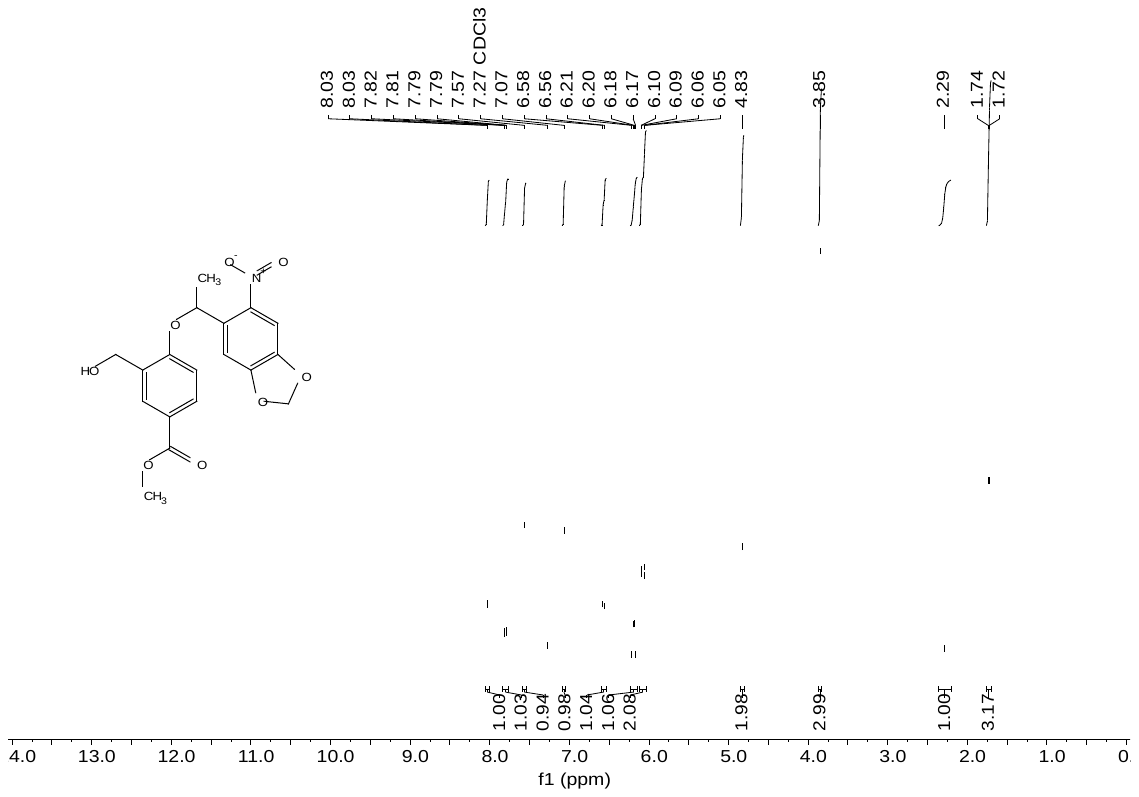

**1c:** ^13^C NMR (151 MHz, Chloroform-*d*)

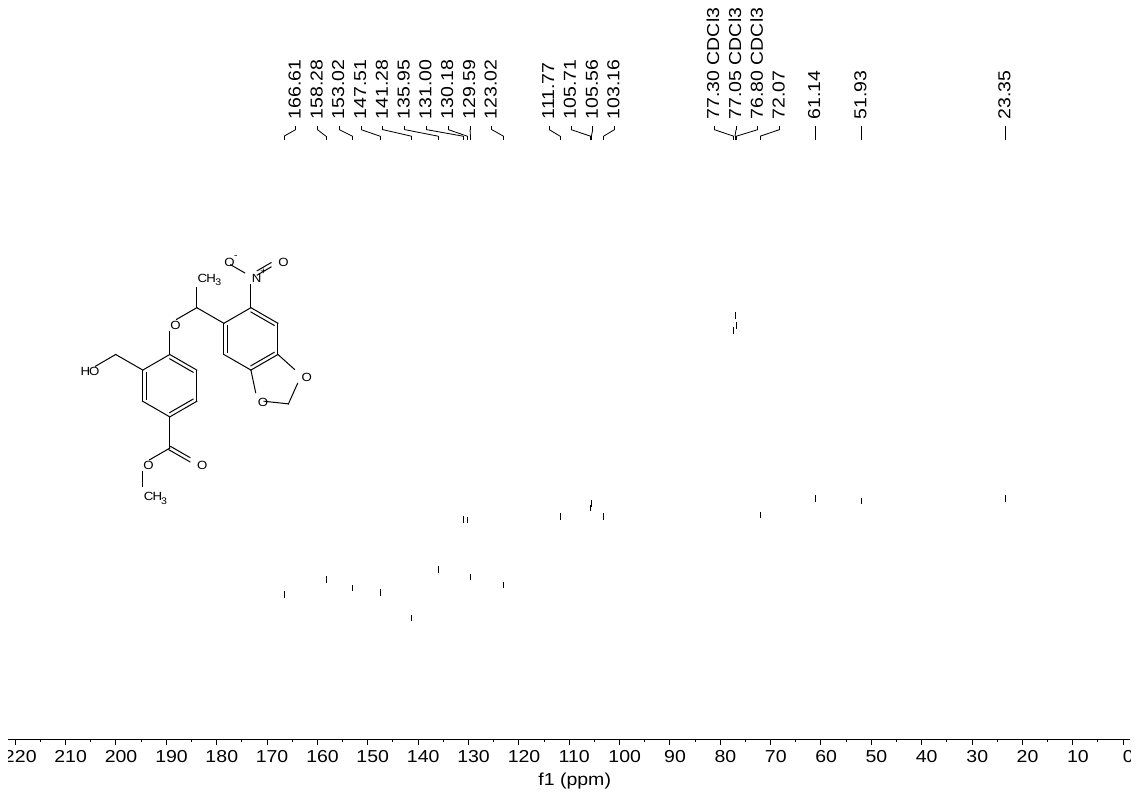

**1d:** ^1^H NMR (400 MHz, Chloroform-d)

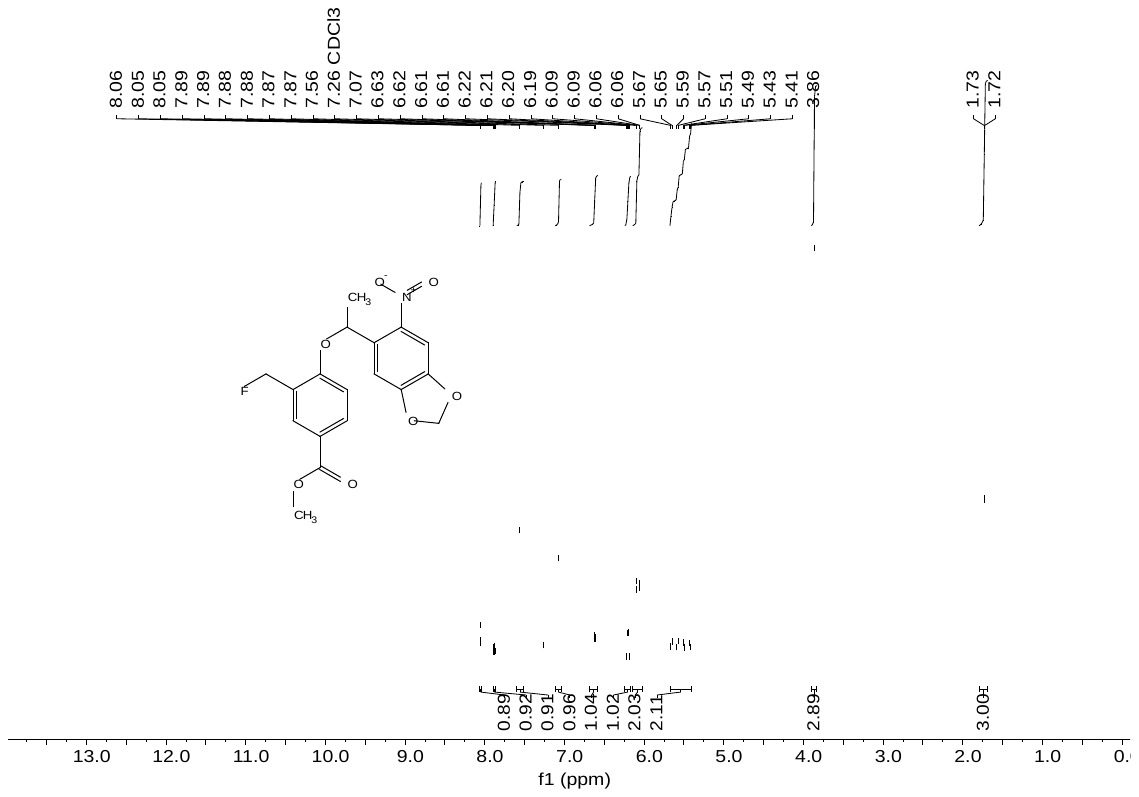

**1d:** ^13^C NMR (151 MHz, Chloroform-*d*)

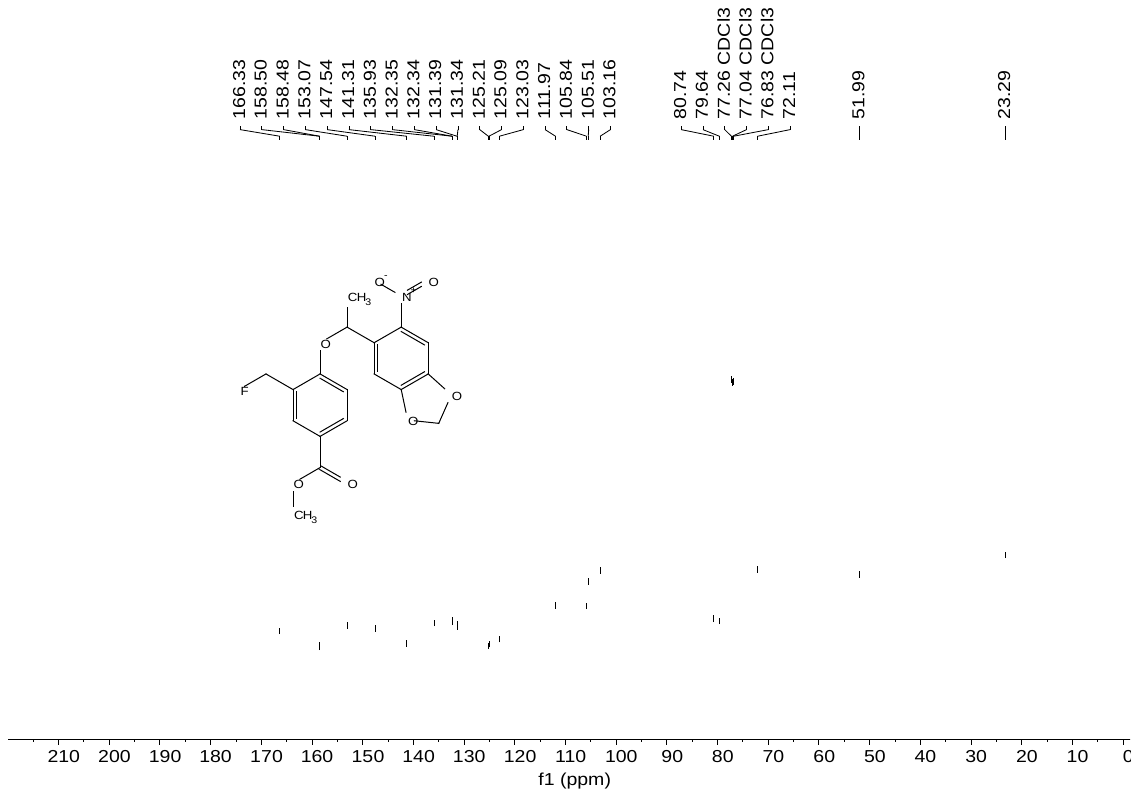

**1f:** ^1^H NMR (400 MHz, Chloroform-d)

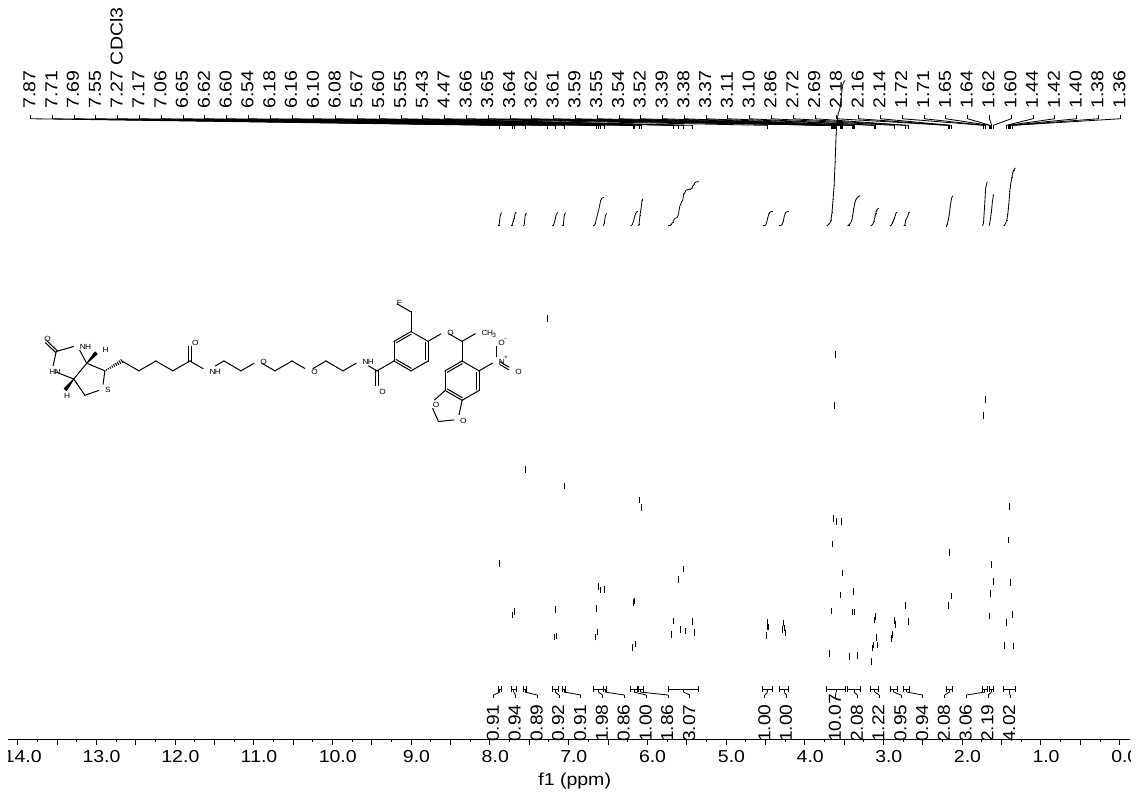

**1f:** ^13^C NMR (151 MHz, Chloroform-*d*)

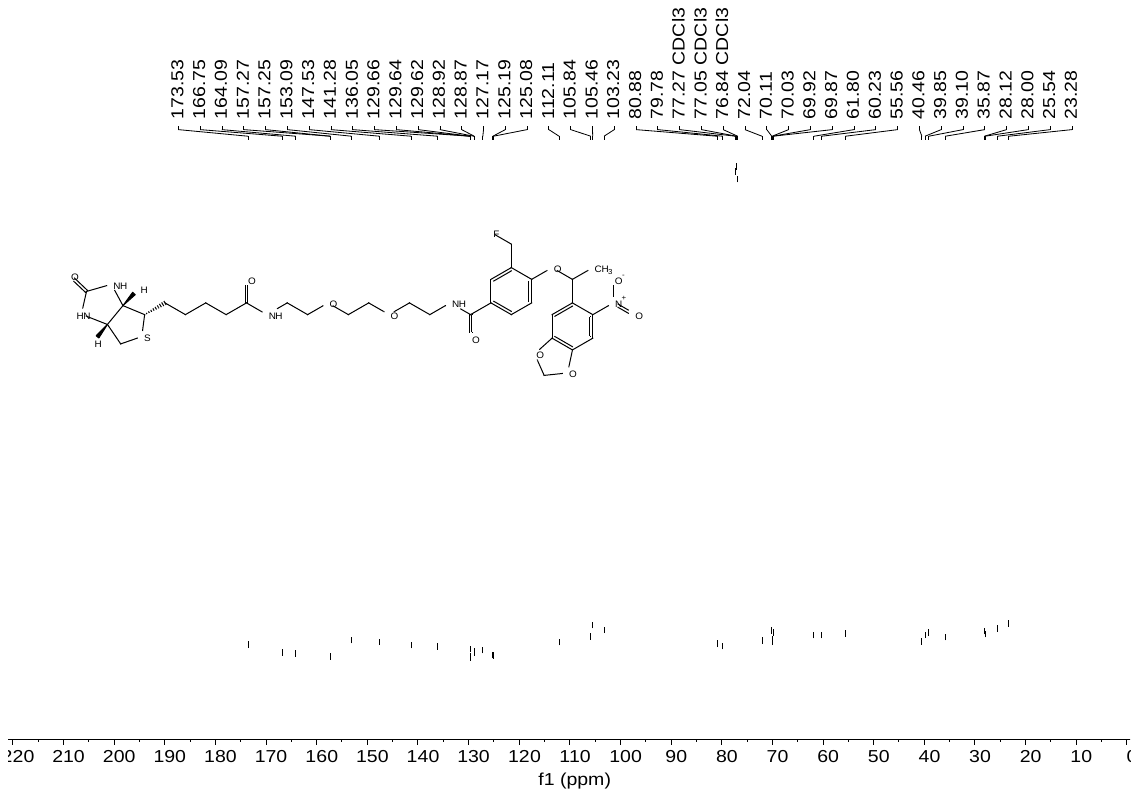

**1f:** ^19^F NMR (471 MHz, Chloroform-*d*)

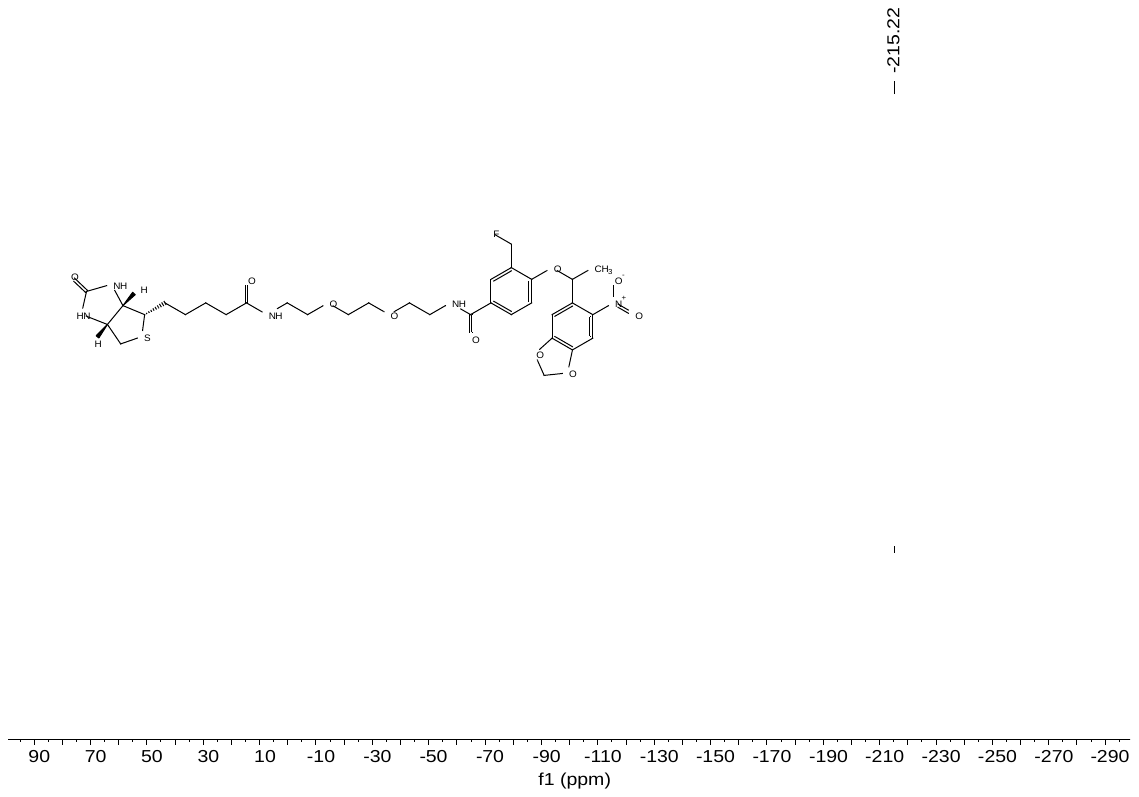

**2b:** ^1^H NMR (400 MHz, Chloroform-d)

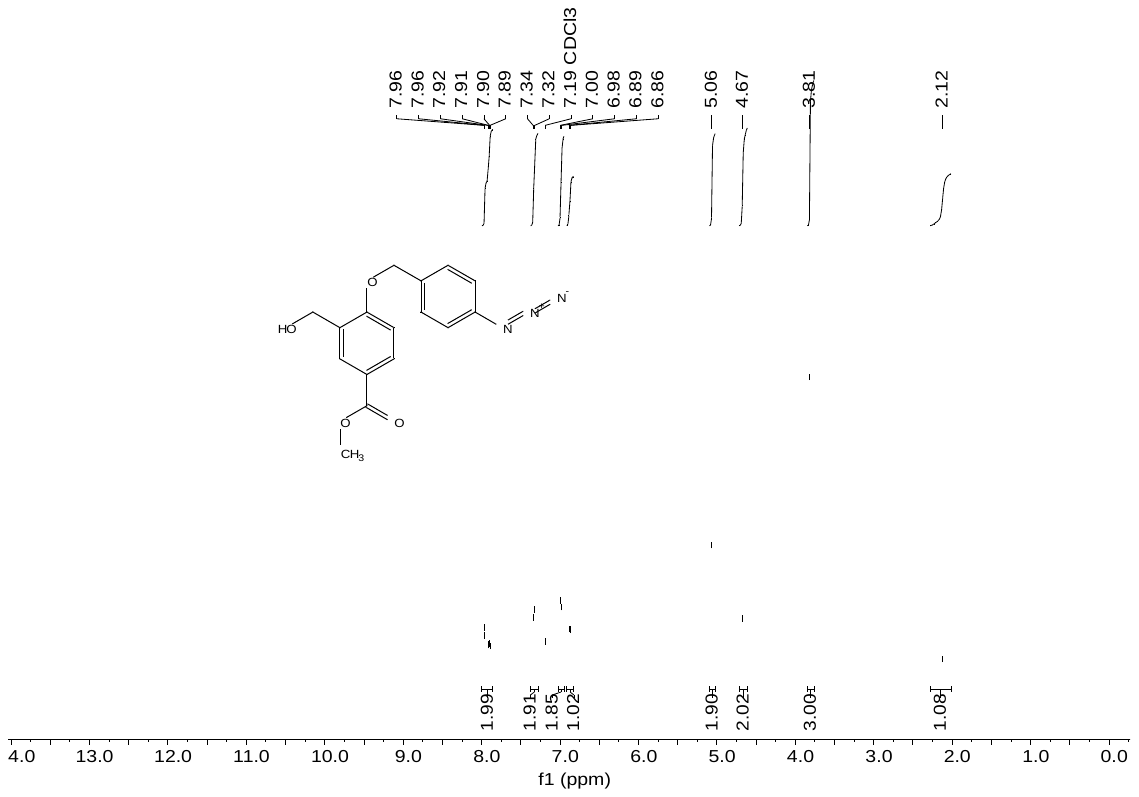

**2b:** ^13^C NMR (151 MHz, Chloroform-*d*)

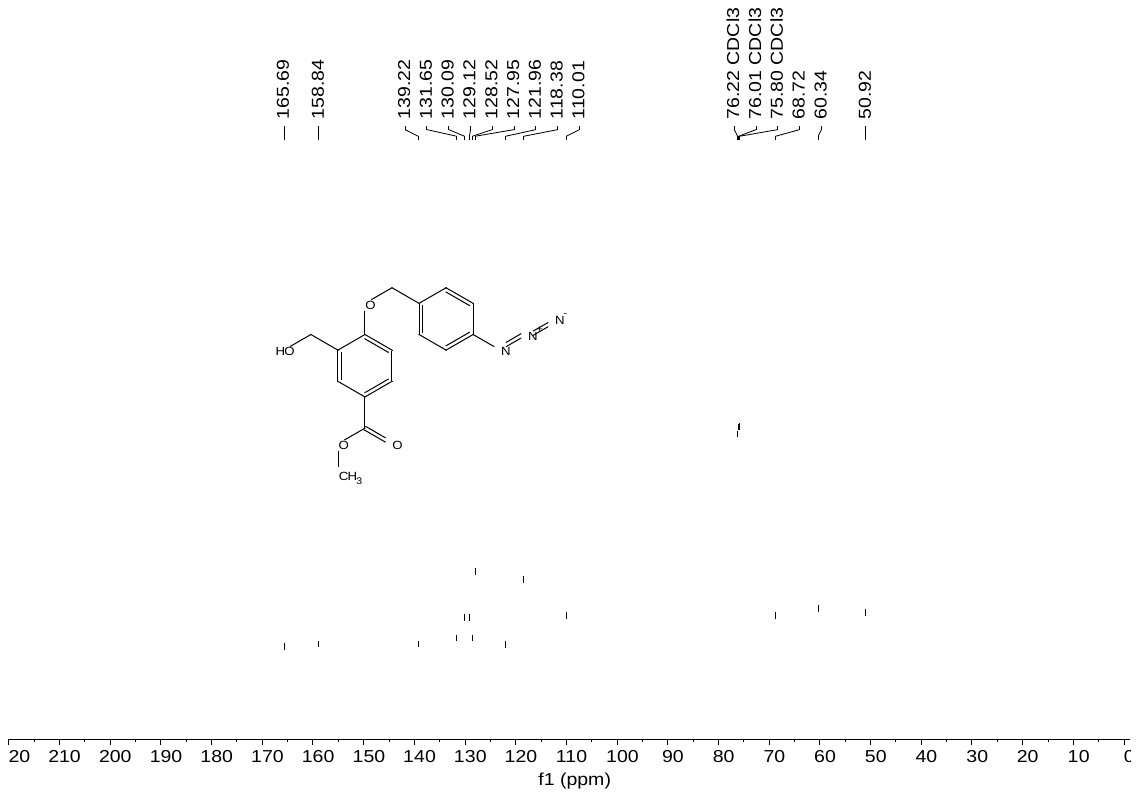

**2c:** ^1^H NMR (400 MHz, Chloroform-d)

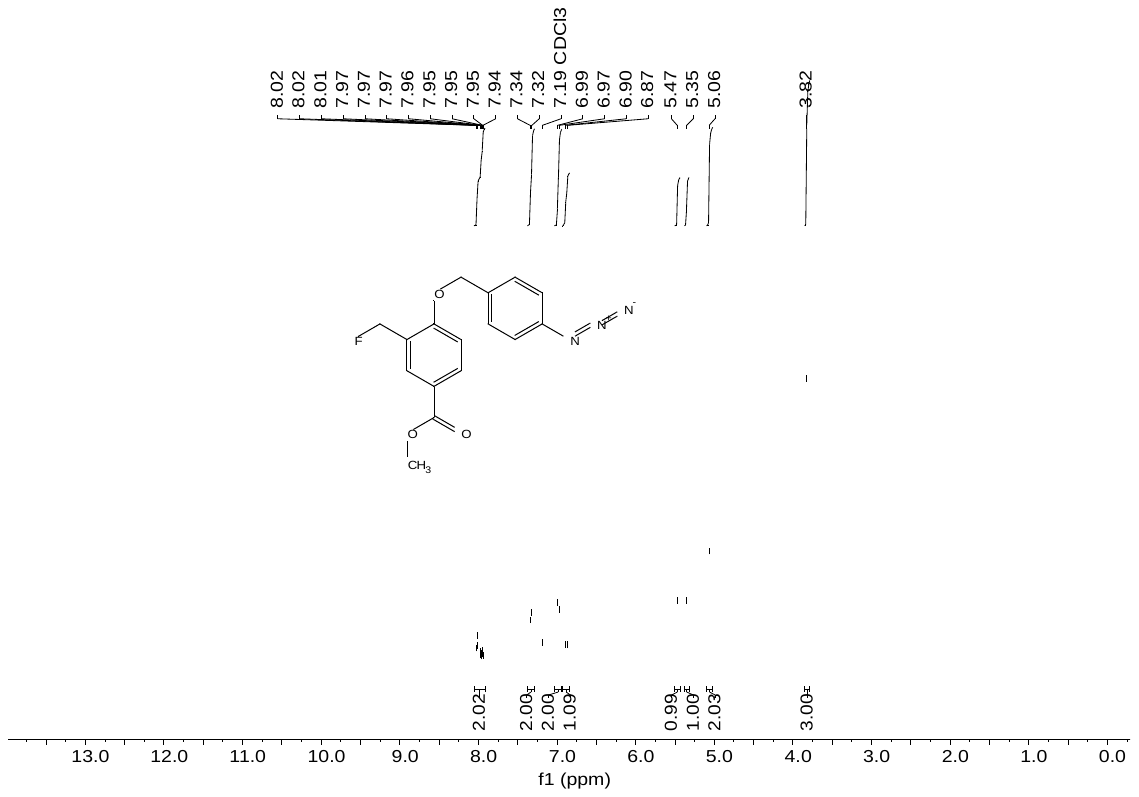

**2c:** ^13^C NMR (151 MHz, Chloroform-*d*)

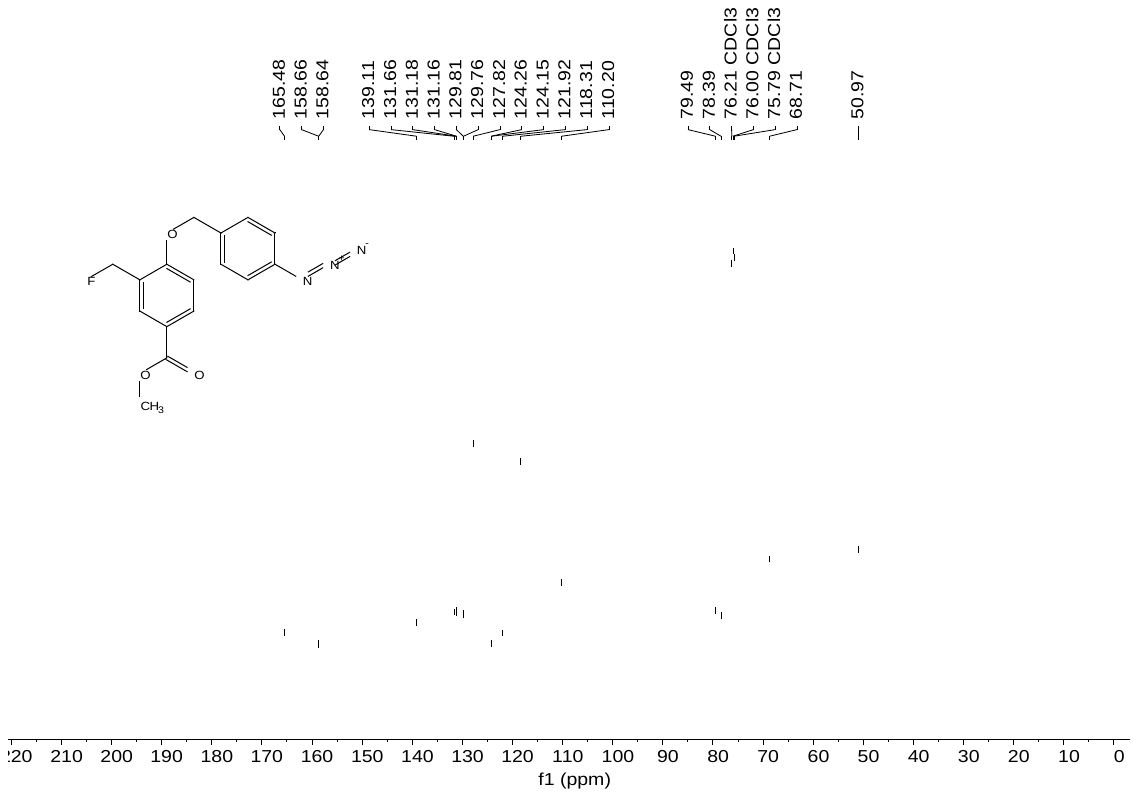

**2e:** ^1^H NMR (400 MHz, Chloroform-d)

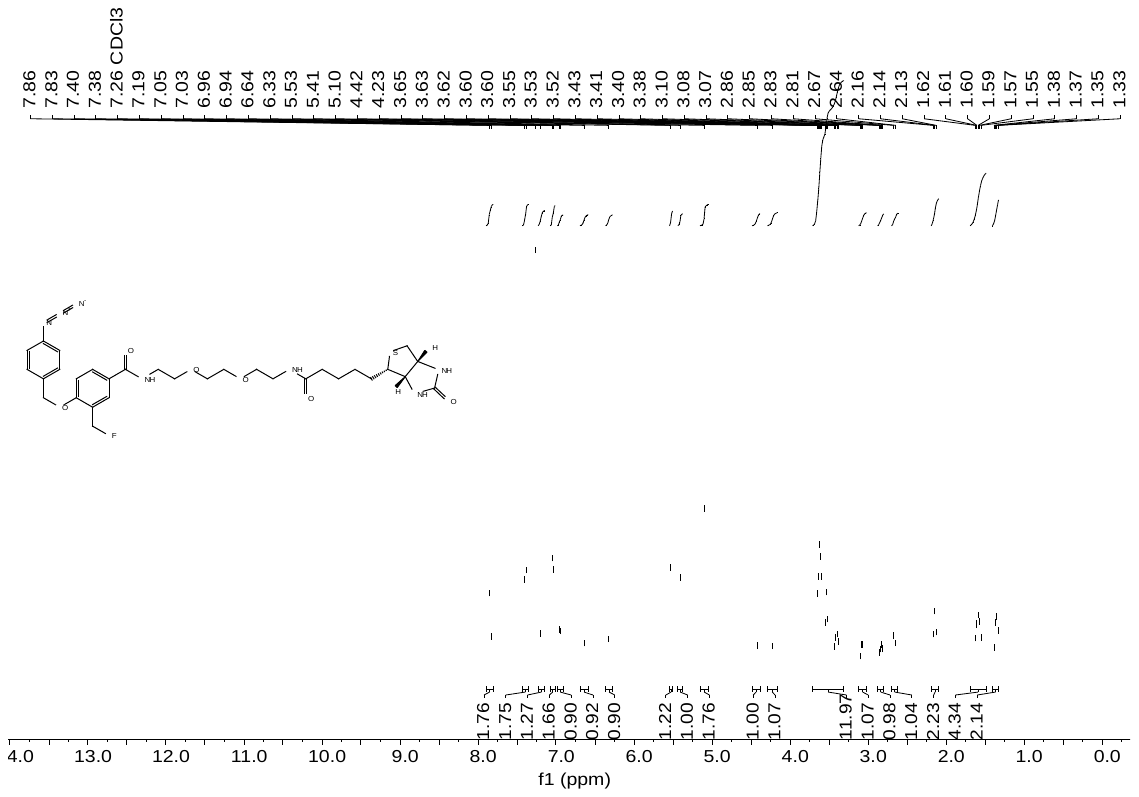

**2e:** ^13^C NMR (151 MHz, Chloroform-*d*)

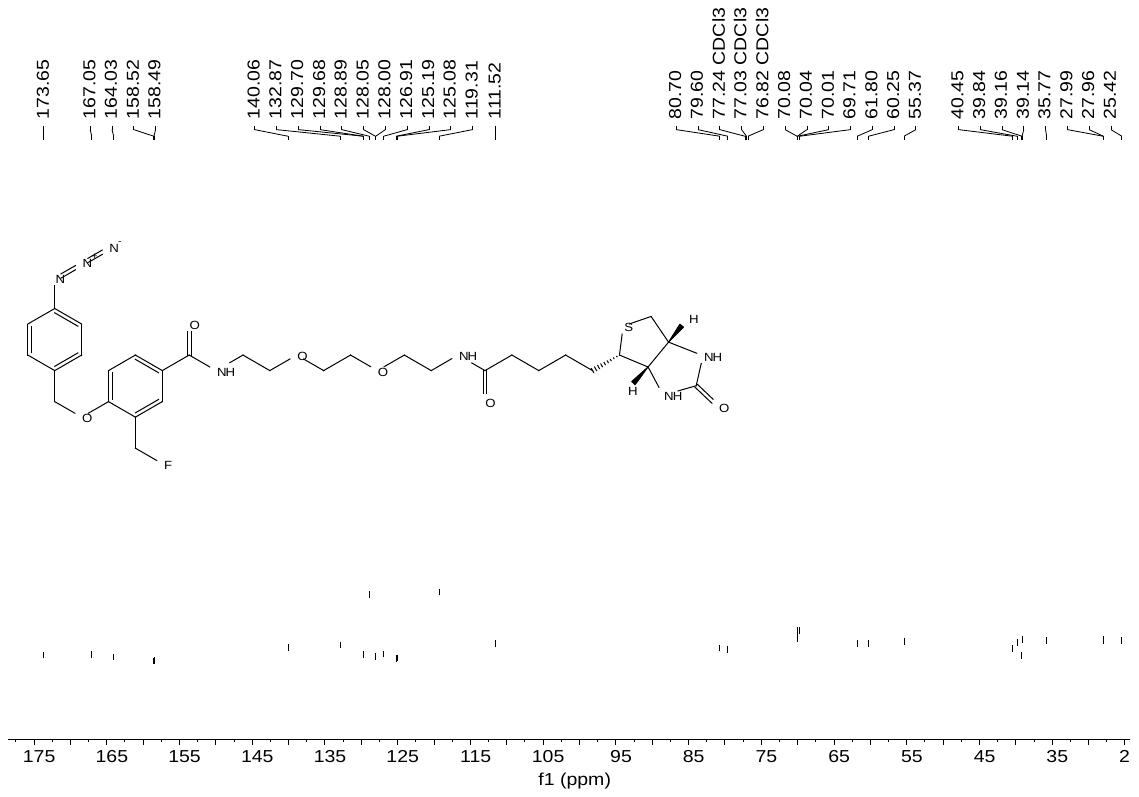

**2e:** ^19^F NMR (471 MHz, Chloroform-*d*)

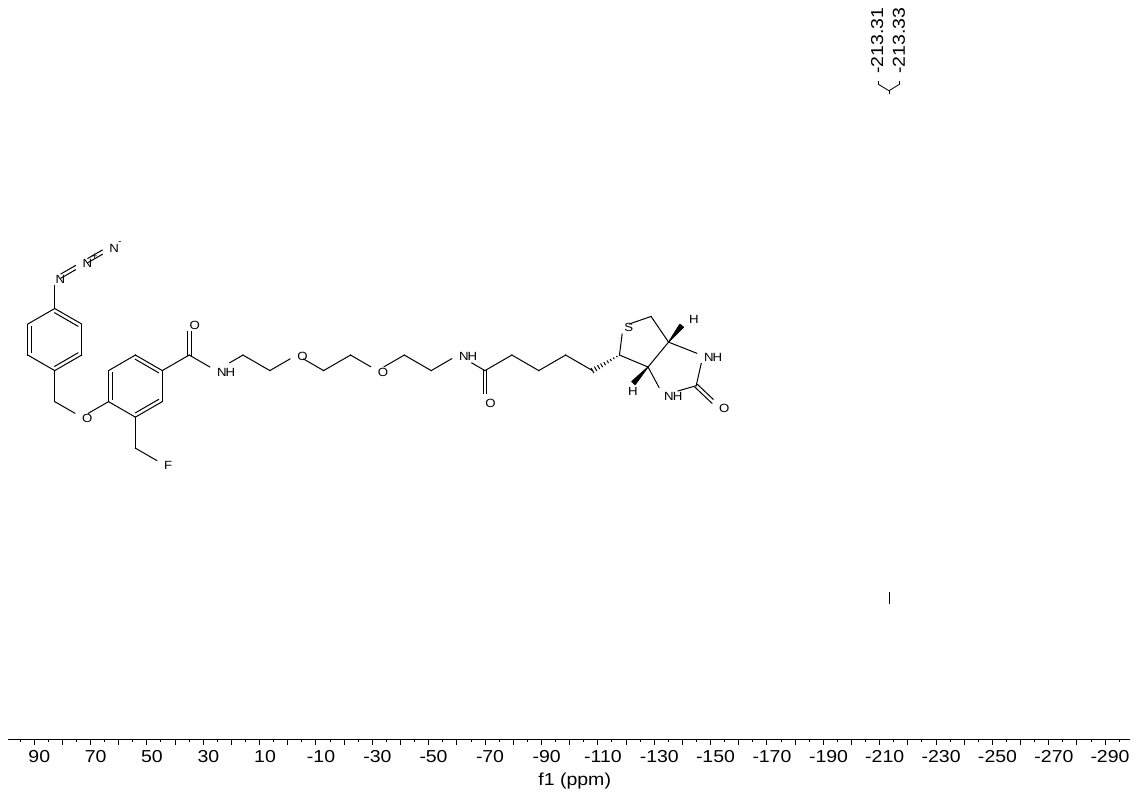

**3a:** ^1^H NMR (400 MHz, Chloroform-d)

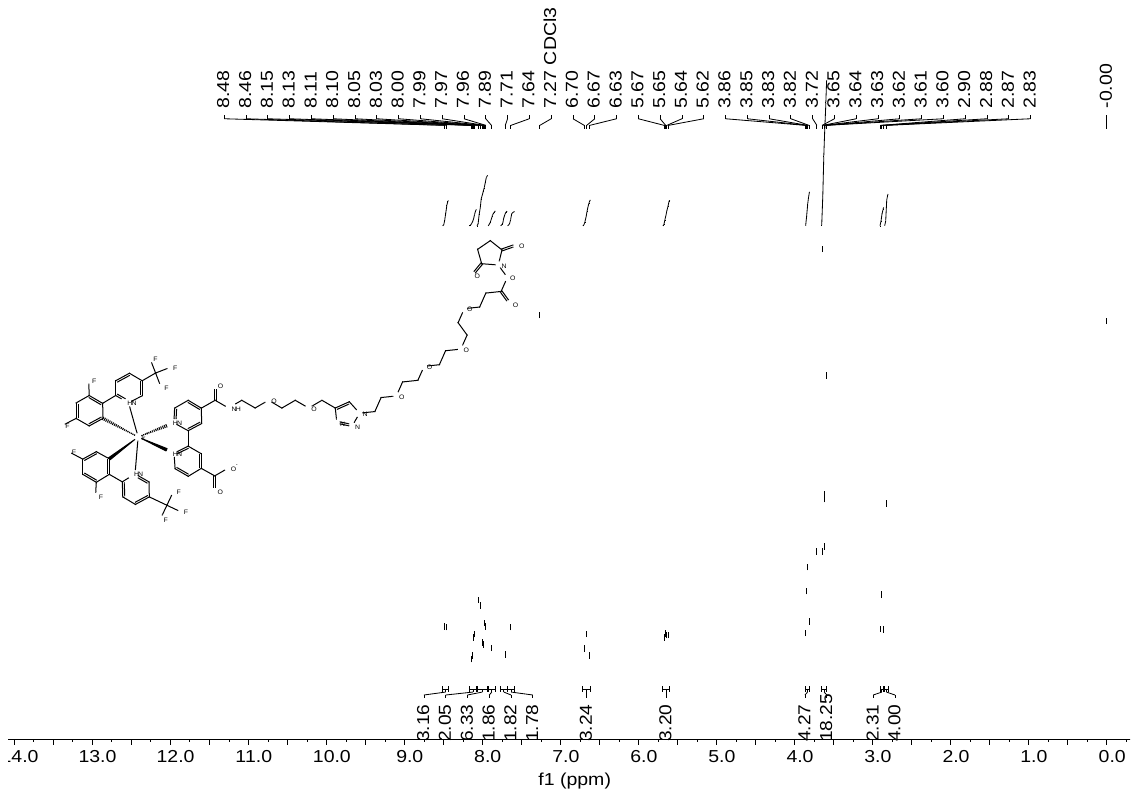

**3a:** ^13^C NMR (151 MHz, Chloroform-*d*)

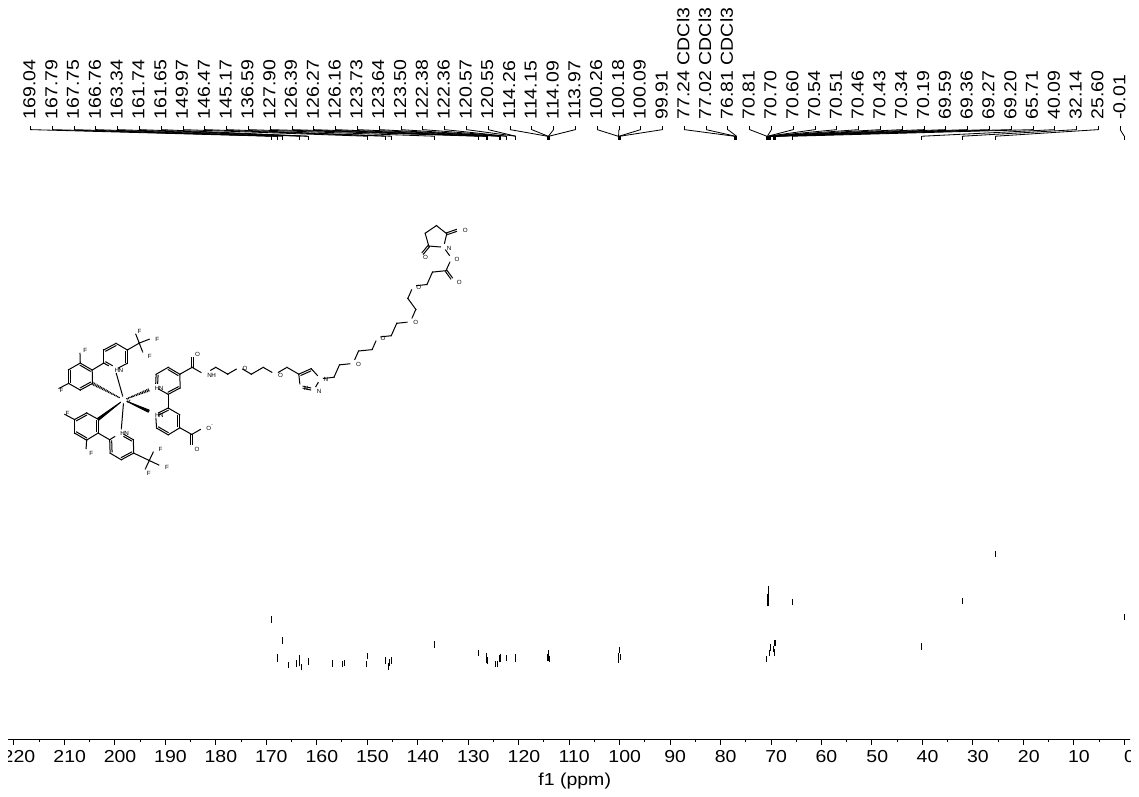
